# Universal architectural concepts underlying protein folding patterns

**DOI:** 10.1101/480194

**Authors:** Arthur M. Lesk, Ramanan Subramanian, Lloyd Allison, David Abramson, Peter J. Stuckey, Maria Garcia de la Banda, Arun S. Konagurthu

## Abstract

What is the architectural ‘basis set’ of the observed universe of protein structures? Using information-theoretic inference, we answer this question with a comprehensive dictionary of 1,493 substructural *concepts*. Each *concept* represents a topologically-conserved assembly of helices and strands that make contact. Any protein structure can be dissected into instances of concepts from this dictionary. We dissected the world-wide protein data bank and completely inventoried all concept instances. This yields an unprecedented source of biological insights. These include: correlations between concepts and catalytic activities or binding sites, useful for rational drug design; local amino-acid sequence–structure correlations, useful for *ab initio* structure prediction methods; and information supporting the recognition and exploration of evolutionary relationships, useful for structural studies. An interactive site, Proçodic, at http://lcb.infotech.monash.edu.au/prosodic (click) provides access to and navigation of the entire dictionary of concepts, and all associated information.

## INTRODUCTION

The polypeptide chains of amino acids (primary structure) in most proteins fold into helices and strands of sheet (secondary structure), which in turn assemble to give proteins their intricate three-dimensional shapes and folding patterns (tertiary structure). Experimental methods have already provided over 140,000 entries in the world-wide Protein Data Bank (`wwPDB`), containing the three-dimensional coordinates of proteins and protein-nucleic acid complexes from a wide range of species.

Unravelling protein architecture and discovering the relationship among these three major levels of structural description provides the key to understanding how proteins function, how their 3D folding patterns form, and how they evolve (1). Investigations of protein folding patterns have revealed recurrent themes at all structural levels (2–8), which form the basis for widely-used hierarchical classifications of protein structures (9–11). Nevertheless, many aspects of the relationships across structural levels have remained unresolved.

Chothia and Lesk (6) introduced the idea of a *core* of the folding patterns of homologous proteins. This core comprises a maximal set of secondary structural elements that assemble in a common 3D topology, while withstanding a certain amount of distortion. The parts outside the core are structurally more variable. Many related proteins contain some but not all of the same common substructures that form their cores. Therefore, it is of crucial interest to discover the nature of the substructures that contribute to the cores of protein families. Some of these are *supersecondary structures* – small conserved combinations of *successive* elements of secondary structure, such as the *β*-*α*-*β* subunit. Supersecondary structures recur within many protein folds, and can be shared even by unrelated proteins. For example, the *β*-*α*-*β* subunit appears in NAD-binding domains, in TIM barrels, and in many other proteins.

Early definitions of supersecondary structures relied strongly on experts spotting and naming them (4, 12). With the steady growth of the `wwPDB`, several methods have been developed to identify automatically, with varying operational definitions, a *library* of substructures that form what can be considered as the 3D building blocks of protein structures (8, 13–25). However, these approaches have yielded limited libraries containing mostly short oligopeptide fragments, or assemblies of typically 2 to 4 secondary structural elements. It has been a challenge so far to go further than that and dissect protein structures into a more complete set that includes *larger* conserved substructures. Apart from the enormous computational challenge this problem poses, the attempts made so far have lacked a statistically-rigorous framework in which to describe, compute, identify and resolve a dictionary of conserved assemblies of secondary structures.

Here we address this problem and unravel protein folding patterns into a universal dictionary of architectural building blocks (*concepts*) containing topologically-conserved assemblies of helices and strands that make contact. This dictionary advances the current knowledge of these conserved patterns significantly beyond the classical supersecondary structures and other known patterns. Our approach relies on a rigorous information-theoretic framework that allows the inference of a dictionary that (a) avoids overfitting (i.e., inferring a dictionary that is more complex than necessary to explain the observed folding patterns) and (b) achieves an objective trade-off between the descriptive complexity of concepts in the dictionary and their fidelity (i.e., the amount of compression) gained when explaining the observed protein folding patterns. Thus, this work presents the ‘basis set’ of concepts underlying all observed protein folding patterns, and allows for any protein chain to be decomposed optimally into parts corresponding to concepts from this set. We put together a comprehensive online resource, Proçodic (click), that can contribute to: understanding the fundamental principles of protein structure, correlating concepts with ligand binding sites to suggest function, and applying sequence conservation observed within concepts to protein structure prediction.

## RESULTS

### Automatic identification of a dictionary of substructural concepts

This work uses the concise *tableau* representation of protein folding patterns introduced by Lesk (26), which is based on the idea that the essence of a protein folding pattern is captured by the order, contacts and geometry of the assembly of secondary structural elements along the amino-acid chain. A tableau corresponds to the 3D structure of a single protein domain (or sometimes chain), and has the form of a symmetric matrix (Fig. 1(a,c); **Supplementary §S1**). Importantly, in this representation supersecondary structures can be defined in a compact and computable way as subtableaux containing two or more *successive* secondary structure elements in contact (Fig. 1(d-e)).

**Fig. 1.**
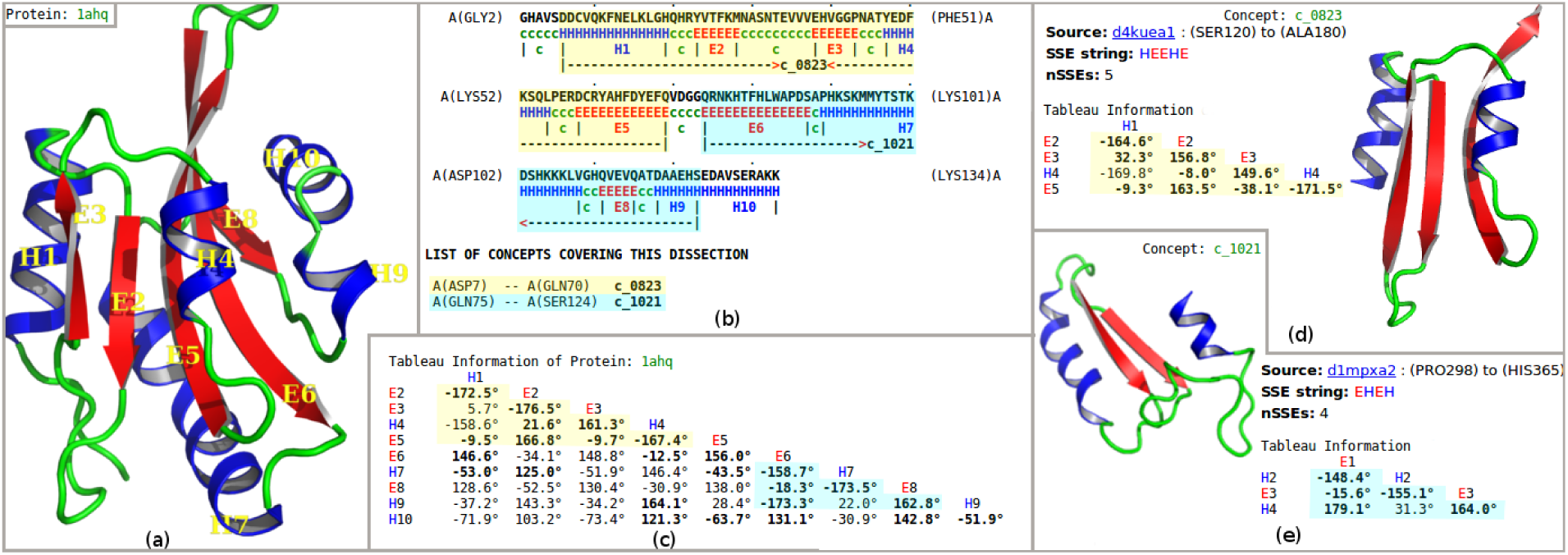
(a) Secondary-structural cartoon representation of the crystal structure of the Actin-binding protein actophorin from *Acanthamoeba* (1AHQ) (32). (b) Secondary structural assignment (using SST (33); H = helix, E = strand of sheet) and the optimal dissection of the protein chain into non-overlapping regions, using the inferred concept dictionary. This information is shown with reference to the amino-acid sequence information in a marked-up format: the dissection of 1AHQ uses *concepts* (see text) `c_0823` (highlighted in yellow) and `c_1021` (highlighted in blue). (c) Tableau representation of the folding pattern of 1AHQ. The highlighted subtableaux correspond to concepts `c_0823` and `c_1021`. Here, only the lower-triangle part of the tableau information is shown because the full tableau is a symmetric matrix. The rows and columns are indexed by secondary structure elements in order of appearance in the polypeptide chain. Off-diagonal elements record the angles between pairs of secondary structural elements; boldface indicates that there is a contact between the corresponding pair of secondary structural elements. (d-e) The concepts `c_0823` and `c_1021` are shown, together with their *archetypal* tableaux and corresponding secondary structural representation.

We have constructed the universal dictionary reported here using our recently-developed method to infer, automatically, conserved assemblies of secondary structural elements within *any* given source collection of tableaux (27). The idea of a concept is constrained by the requirement that every secondary structural element in the concept must be in contact with at least one other secondary-structure element in that concept. Our concept inference approach (27) is based on the powerful minimum message length criterion for statistical inductive inference (28–30) and lossless data compression (**Supplementary §S2**). We have applied this method to compress the source collection of Astral SCOP domains (9, 10, 31) (**Supplementary §S1**). This has allowed us to infer a dictionary of 1,493 substructural concepts that *most concisely* and *losslessly* describes the entire source collection, and does so without any prior knowledge or preconceived notions regarding these recurrent substructures.

The total computational effort required to identify this dictionary is equivalent to about 7 years of runtime on a modern computer. Therefore, we parallelised our method and ran it on a high-performance computing cluster using 240 cores to identify the Proçodic dictionary in 14 days (**Supplementary §S2**).

### Proçodic: The dictionary of inferred concepts

Each of the 1,493 concepts in the dictionary is designated by an identifier of the form ‘`c_`’ followed by 4 digits: `c_0001`—`c_1493`. This order follows (1) the decreasing length in the number of secondary structural elements (nSSEs) defining each concept, and the lexicographic order of their secondary structural strings, where we represent any helix by ‘`H`’ and any strand by ‘`E`’.

Fig. 2 shows the top 100 concepts in the dictionary. The largest concept (`c_0001`) contains 28 secondary structural elements. The smallest concepts (`c_1441`—`c_1493`) – not shown in Fig. 2 – contain only 2 elements. (Note that a single helix or a single strand/extended region is not considered here as a concept.) The distribution of inferred concept sizes is shown in Fig. 3(a): 9 concepts (`c_0001`—`c_0009`) are composed of an assembly of ≥ 20 secondary structural elements (SSEs), 48 concepts (`c_0010`— `c_0057`) have between 15 and 19 SSEs, 217 concepts (`c_0058`—`c_0274`) contain between 10 and 14 SSEs, 217 concepts (`c_0058`—`c_0274`) contain between 10 and 14 SSEs, and 368 concepts (`c_0275`— `c_0642`) contain between 9 and 6 SSEs. The remaining concepts contain between 5 and 2 SSEs. The median concept size is 5 SSEs.

**Fig. 2.**
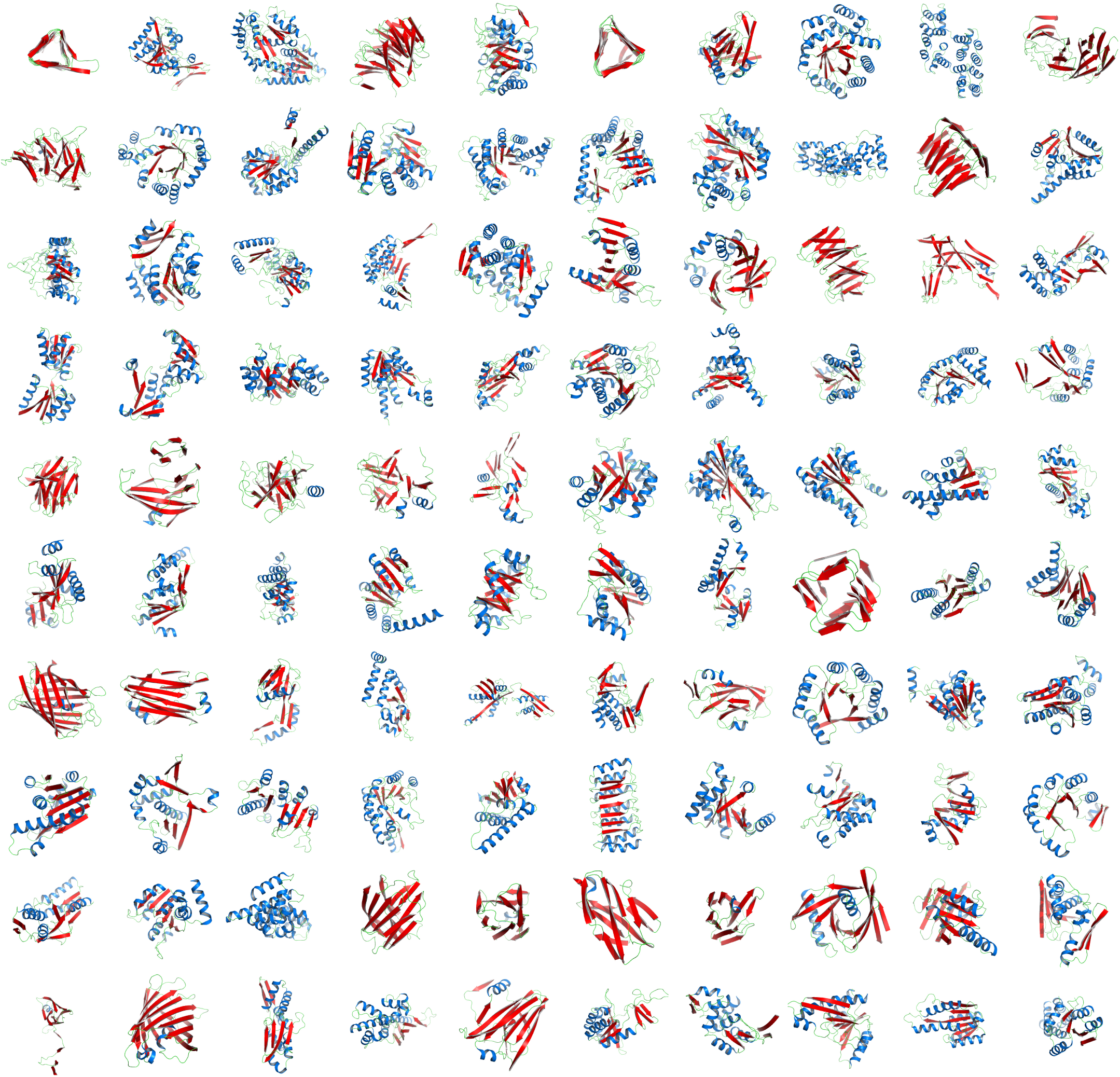
Representative structural cartoons of the top 100 concepts from the inferred dictionary containing 1,493 concepts, ranked in decreasing order of number of secondary-structure elements (row-wise top-left to bottom-right: `c_0001` to `c_0100`). Strands of sheet are shown in Red; helices in Blue. (See website for the full interactive listing.) The inference of the whole dictionary is automatic without any prior knowledge or preconceived notions of these recurrent themes. The inferred concepts subsume known patterns, for example: ‘*α*-*β* Barrel’ (`c_0005`), ‘Armadillo repeat’ (`c_0083`), ‘*β* Barrel’ (`c_0061`), ‘*β* Propeller’ (`c_0004`), ‘Icosahedral (Virus)’ (`c_0067`), Immunoglobulin (`c_0062`), ‘Jellyroll architecture’ (`c_0084`), ‘Left-handed *β*-Helix’ (`c_0001`), ‘Leucine-rich repeat’ (`c_0076`), ‘Right-handed quadrilateral *β*-Helix’ (`c_0058`) ‘NAD-binding domain’ (`c_0002`), ‘TIM barrel’ (`c_0008`) etc. Other classical supersecondary structures not shown in this figure such as *β*-hairpin (`c_1442`), *α*-hairpin (`c_1484`), *β*-*α*-*β* unit (`c_1240`) appear lower down in the dictionary of concepts, ordered from largest to smallest. See text (Page 5) where classical supersecondary structural motifs are discussed.

**Fig. 3.**
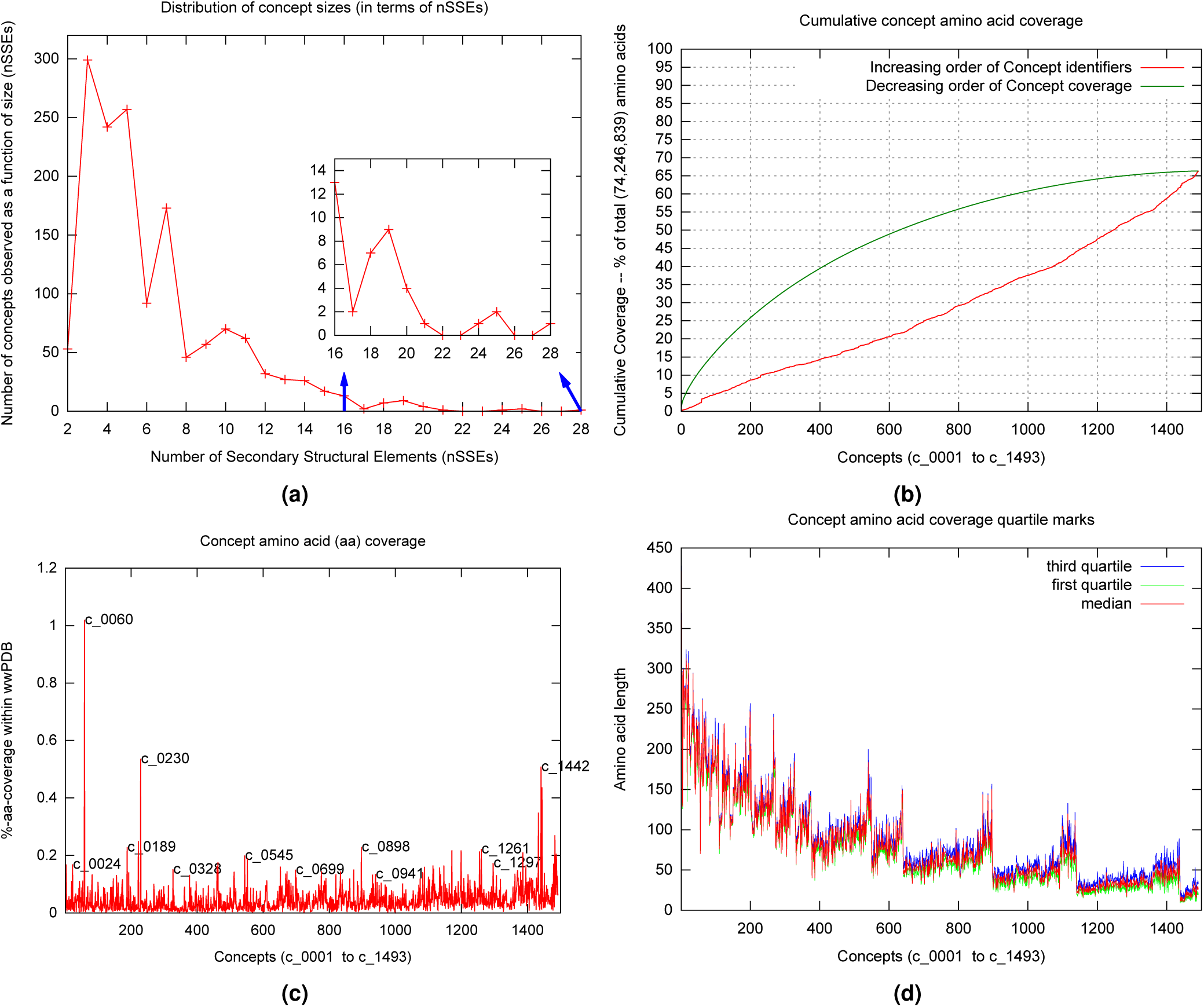
(a) Distribution of concept lengths in terms of the number of secondary structural elements (nSSEs) they contain. The smallest concepts have 2 secondary structural elements; the largest has 28. (b) Cumulative amino acid coverage of concepts (as a percentage of the total 74,246,839 number of residues) after dissecting 275,014 protein chains using the inferred dictionary. The green curve gives the distribution in the decreasing order of individual concept amino acid coverage – i.e., the concept with largest coverage is listed first, that with second largest coverage is listed second, and so on. The red curve gives the same cumulative distribution in the serial order of concept identifiers – i.e., concept `c_0001` is shown first, concept `c_0002` second and so on. (c) Individual concept amino acid coverage (as a percentage of the total 74,246,836 residues) in the serial order of concept identifiers (with some concepts highlighted). (d) Dissecting the protein chains from the `wwPDB` allows us to catalogue the regions where each concept is used. Underlying each concept usage is an amino acid sequence of variable length (although the associated strings corresponding to the types and order of secondary structural elements match exactly). This graph plots the first, second and third quartile points in the distribution of amino acid lengths for each concept’s set of usages in the `wwPDB`. Concepts are listed in the decreasing order of lengths, followed by the lexicographic order of their secondary structural strings. Since the average strand of sheet (denoted as ‘E’) has fewer amino acids than the average helix (denoted as ‘H’), the lexicographic order creates in the plot the observed piecewise increasing trend among concepts with same number of secondary structural elements.

The complete inferred dictionary is available via the interactive website Proçodic (for Protein Concept dictionary – the cedilla allows the pronunciation as ‘prosodic’) at http://lcb.infotech.monash.edu.au/prosodic. As discussed below, this site allows the exploration of any structure the user provides as input, or specific concepts that are of motivating focus for the user, including: the usages of concepts in other structures, both homologous and non-homologous; or the inspection of frequently occurring keywords within the ‘KEYWDS’ records and the ligand-binding information from the ‘HETATM’ records extracted from the source `wwPDB` coordinate files (see **Discussion**).

### Our dictionary subsumes known supersecondary structural motifs

Our automatically identified dictionary includes many concepts that match the known repertoire of supersecondary structural motifs (34). Matched motifs involving assemblies of a small number of helices and strands include: antiparallel (`c_1442`) and parallel (`c_1443`) *β*-*β* assemblies, *α*-*α* hairpin (`c_1484`) *α*-*β*/*β*-*α* assembly, (`c_1459`/`c_1472`), basic helix-loop-helix (`c_1351`), *β*-*α*-*β* motif (`c_1240`), EF-Hand (`c_1342`, `c_1491`), *f*-motif (`c_1178`), helix-turn-helix motifs (`c_0826` – winged type I, `c_0870` – winged type II, `c_1373` – plain), four-helix bundle (`c_1101` – type I, `c_1117` – type II), *β*-meander (`c_1187`), Greek key (`c_0964`), Zinc finger (`c_1230`), helix-hairpin-helix motif (`c_1068`), *β*-sandwich (`c_0390`), and *αβ*-sandwich (`c_0603`) among others.

Our dictionary also includes larger assemblies of helices and strands that match known *repeating* structural motifs. These include: three-sided left-handed *β*-helix (`c_0001`, `c_0380`), three-sided right-handed *β*-helix (`c_0388`), right-handed quadrilateral *β*-helix (`c_0058`), ankyrin repeat (`c_0370`, `c_0632`), armadillo repeat (`c_0083`, `c_0888`), kelch repeat (`c_0395`), *α*-solenoid (`c_0270`, `c_0271`), and Leucine rich repeat (`c_0076`) among others.

To understand the relationships between different concepts, we have clustered hierarchically the 1,493 Proçodic concepts. Since each concept archetype defines a (sub)tableau derived from a tableau of the domain in the source collection, we start by inferring the dictionary of *meta-concepts* that best explains all the Proçodic concept tableaux. This is achieved by using exactly the same unsupervised inference methodology that was used to infer Proçodic concepts. That is, we now treat the tableaux representing 1,493 archetypes from our inferred prosodic concept dictionary as the source collection, and rerun our inference method (**Supplementary §S2**). This yielded 34 meta-concepts that dissect (i.e., best explain) the inferred 1,493 concepts. The text file containing these meta-concepts, along with the corresponding list of Proçodic concepts that use each meta-concept within their dissections, is available in the supporting data file: `metaConceptsAndUsageList.txt` (click).

Once the meta-concepts are obtained, we dissect each of the 1,493 concepts using these meta-concepts. This permits a 34-dimensional feature vector representation of concepts, where each vector-component denotes the number of times the corresponding meta-concept is used in that concept dissection. We note that this representation is similar to the bag-of-words model (35) used in information retrieval and natural language processing. Using this feature vector representation, the 1,493 Proçodic concepts are clustered hierarchically by:

1. Constructing a 1, 493 *×* 1, 493 similarity matrix that compares all pairs of these 34-dimensional vectors, and
2. Using the resultant similarity matrix to cluster all Proçodic concepts hierarchically, based on the unweighted pair-groups method using arithmetic averages (36).

This procedure yielded a hierarchical tree of concept relationships, available in interactive format from: `prosodicConceptClustering.html` (click). This tree reveals similarities that are also detectable by examining the concept archetypes, their usages and keywords. For example, `c_0009` and `c_0018` are both helical bundles related to the architecture of Annexin proteins, with `c_0009` having one extra helix compared to `c_0018`. Another example is the cluster containing `c_0001`, `c_0006`, `c_0113`, and `c_0380`, where all represent left-handed *β*-helical motifs composed of 28, 20, 12 and 7 *β*-strands, respectively.

Although our average concept archetype is significantly smaller (with 47.6% of the number of SSEs) than its source protein domain, several concepts inferred in our dictionary describe conserved folding patterns at the level of domains. These include: NAD-binding domain (`c_0173`), *β*-grasp fold (*e.g.* `c_729`*), β*-propeller (`c_0382`), Swiss/Jelly roll fold (`c_0406`), Ferredoxin (plait) fold (`c_0581`), TIM barrel (`c_0008`), Immunoglobulin fold (`c_0118`, `c_0121`), Ubiquitin roll (`c_0737`), and large *β*-barrel (`c_0061`).

These results show that our dictionary encompasses a broader set of substructural invariants across the protein folding space than previous studies achieved (see section ‘Comparison with previous related work’). This advantage is mainly due to our use of tableaux to capture concisely the essence of protein folding patterns, and our use of the information-theoretic criterion of minimum message length to yield an objective dictionary complexity-versus-fidelity trade-off.

### Dissection of wwPDB and coverage of concepts across the protein folding space

The methods used for this work also permit the optimal *dissection*, within seconds, of any protein chain into non-overlapping regions that are explained (compressed) using the concepts from the inferred dictionary. Regions not assigned to any dictionary concept (notionally designated to the *null* concept, `c_0000`) remain uncompressed. These include the small set of proteins that have no secondary structure, for instance wheat-germ agglutinin (9WGA). Fig. 1 shows an example of the dissection of the crystal structure of the Actin-binding protein actophorin from *Acanthamoeba* (1AHQ). (See Proçodic website to dissect any protein structure of interest.)

We have dissected the entire `wwPDB`, which at the time of calculation resulted in tableaux corresponding to 275,014 protein chains containing 74,246,839 amino acid residues overall. (Note that the dictionary was constructed using an *unbiased* set of domains from ASTRAL; but the subsequent dissection of the entire `wwPDB` reflects the biases in the distribution of protein folding patterns in the full database.) The usages of the resulting concepts cover regions within proteins that account for 66.35% (49,262,577) of the total (74,246,839) amino acids in the `wwPDB` protein chains we dissected. The remaining 33.65% is dominated by single secondary structural elements, plus loops between successive concept assignments along a dissected chain. Fig. 3(b-d) show the distributions of amino acid coverage of concept usages within the `wwPDB`. Concept `c_0060` (click) has the largest coverage in terms of the number of amino acids its usages cover. This concept is composed of 14 secondary structural elements (SSE string: EEEEHHEEEEHHEE) assembling into a four layer architecture, with its core containing two layers of closely-packed five-stranded *β*-sheets (37) that are sandwiched between two outer layers, containing two *α*-helices each. In total, this concept was used within 3,892 protein chains, with a median value of amino acid coverage equal to 194 residues. Examination of these usages reveals that they come from the protein chains of 285 proteasome complexes. At the other extreme is concept `c_0568` (click), which has the smallest amino acid coverage: 561 residues over 13 protein chains related to plant and bacterial Ferredoxins (38). This concept is composed of 6 secondary structural elements (SSE string: EEHEEE).

Novel insights regarding the concepts can be gained from their usage information. For example, consider the concepts `c_0060` and `c_0568` mentioned above: the concept `c_0060` covers the *β*5 subunit of a recently solved structure of the native human 20S proteasome at 1.8 Å resolution (5LE5) (39). This landmark study revealed a number of functionally important differences with respect to what was known from the previously published 20S proteasome structures. In particular, it identified chloride ions within all active sites, thus significantly revising the description of the proteasome active site, and providing new insights into peptide hydrolysis that underpin the ‘*development of next-generation proteasome-based cancer therapeutics*’ (39). Examination of the usages of `c_0060` within the dissection of 5LE5 (chain Y – *β*5 subunit), reveals that this concept is directly linked to proteolytic active sites (Fig. 4(a)). Analyses of the human-annotated keywords used in the `wwPDB` coordinate files from these usages showed among its top 10 frequently used phrases terms such as ‘Cancer (therapy)’, ‘Drug resistance’ and ‘Bortezomib’ – an anti-cancer drug and the first therapeutic proteasome inhibitor to be used in humans (see Proçodic). This is strong evidence of the concept being linked to a proteolytic active site. A similar examination of the usage instances of the concept `c_0568`, directly links it to the Fe2S2-cluster binding Ferredoxins (see Fig. 4(b)), which mediate electron transfer (40).

**Fig. 4.**
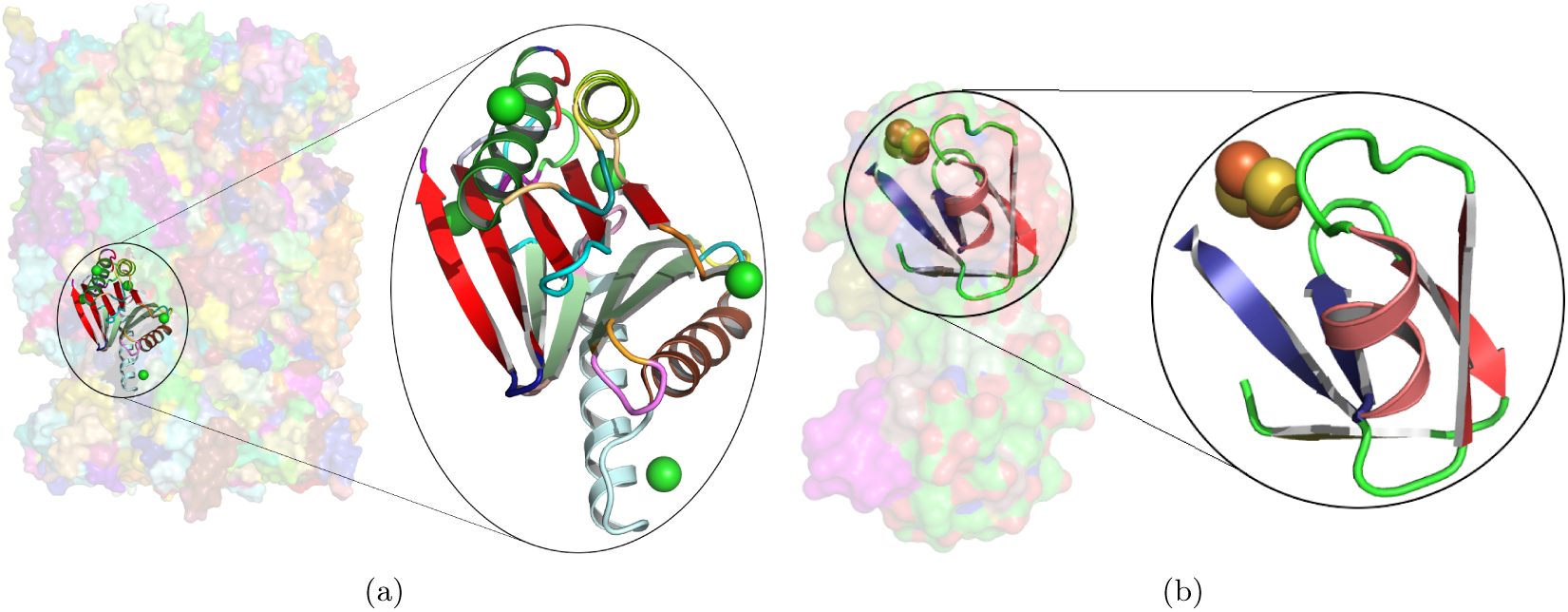
(a) Transparent surface rendering of the native human 20S proteasome at 1.8 Å (5LE5), with the usage of concept `c_0060` in the *β* 5 subunit (chain Y in the amino acid region THR1 to ASN191) shown in1 cartoon. The closeup of this region reveals a chloride ion in all active sites. Chloride ions are known to facilitate a proton shuttle catalytic mechanism (39). (b) Similar rendering as above for the usage of concept `c_0568` in the 2.3 Å Ferredoxin structure from *Mastigocladus laminosus* (3P63 chain A in the amino acid region THR48 to GLU90). The closeup shows the region linked to the Fe_2_S_2_-cluster binding.

### Comparison with previous related work

Many previous studies have attempted to identify a canonical set of recurrent patterns within the structures of proteins. Among the earliest such studies is the seminal work by Unger and colleagues (13), who demonstrated that most hexapeptide fragments in the (then) known proteins are structurally similar to a set of about 100 representative fragment types, using a normalised root mean square devision (RMSD) based clustering methodology. Much subsequent work have followed novel variations of this RMSD-based clustering strategy involving short oligopeptide fragments (of differing length) to produce different fragment libraries (8, 16–19, 41–43). Among other noteworthy approaches to roster oligopeptide fragments are the investigations of Baker and coworkers, who studied the distributions of local structures by clustering short amino acid sequence information (44, 45). Their fragment libraries, together with the inferred local sequence-structure relationships, now underpin successful and popular *ab initio* structure prediction methods (46). Further, Nepomnyachiy et al. (23) recently proposed a pipeline to explore reuse of regions in proteins based on their amino acid sequence relationships. This work reported reuse of sequence segments between 35 and 200 amino acids in length. However, relying on amino acid sequence relationships to identify reuse is rather limiting, as sequences diverge more drastically than structures in evolution (6).

Focusing on the methods that rely on structural information, Grishin and colleagues (20) recently proposed a method to enumerate constructively all idealised *parallel/antiparallel* arrangements of up to 5 SSEs. This work proposed a systematic enumeration of all possible parallel/antiparallel arrangements using a 3D lattice model. This allowed them to model theoretical arrangements of SSEs and use them to search for observed occurrences of each arrangement within the PDB. However, their idealised models are limited to parallel/antiparallel orientations, which poses a considerable restriction in exploring the full set of SSE arrangements observed in the PDB.

Furthermore, two new types of motif libraries have also been recently proposed: the Smotif library (21) and the TERMs library (22). An Smotif is designated by the arrangement of a *pair* of SSEs (of one of the following types: EE, EH, HE, and HH). A library of Smotifs is a collection of such SSE-pairs with different geometries. Their work utilises an RMSD theshold of 2.5Å to cluster 11,068 observed pairs of SSEs from a collection of 1,200 protein structures (i.e., one randomly-chosen protein domain per SCOP fold). These fragments serve in their work as the representatives of the protein structural space. Thus, any consecutive pair of secondary structures within a protein chain is assigned to the closest (based on RMSD) representative Smotif.

Beyond these basic pairwise assemblies identified by the Smotif library, the tertiary motif (TERMs) library (22) was able to find bigger assemblies of short oligopeptide fragments using the following approach. For each amino acid residue *i* in the non-redundant collection of 29,000 residues, a candidate TERM is defined using one or more oligopeptide fragments formed by the union of the residues *i -* 2, *…, i* + 2 together with all penta-peptide regions around residues that form a ‘potential contact’ with the residue *i*. For each candidate TERM, the method finds matching tertiary fragments using an RMSD-based search method. A subset of candidate TERMs is realised by posing it as the classical set cover problem and realising the minimal cover using a greedy approximation method that iteratively identifies the TERMs (based on their coverage) that match proteins in the considered set. This iterative procedure yields about *half a million* (458,251) TERMs. The minimum TERM has 1 oligopeptide fragment containing 5 amino acids, while the maximum TERM has 10 fragments with 52 amino acids. Importantly, an average TERM in their library is composed of 3 oligopeptide fragments covering 19 amino acids (i.e., 6 amino acids per fragment). Furthermore, inspecting the TERMs that cover 50% of their proteins in their considered collection of 29,000 protein structures, we find that each TERM averages 2 fragments with 12 amino acids. Moreover, inspecting the top 24 TERMs (see Fig 2A of (22)), we find many repetitions of short helices and antiparallel strands.

In comparison, our work results in only 1,493 architectural concepts (two orders of magnitude more concise than TERMs), where our minimum concept is composed of 2 SSEs whose median usage in the PDB covers 19 amino acids, while the maximum is composed of 28 SSEs whose median usage covers 171 amino acids. An average prosodic concept in our dictionary is composed of 6 SSEs covering 75 amino acids. Considering the prosodic concepts that cover 50% of the PDB, an average concept has 5 SSEs covering 66 amino acids. Thus, using this framework, our dictionary yields concepts that are a substantially larger than TERMs, and define a significantly more economical dictionary that explains the entire PDB. Moreover, our methodology defines a direct and efficient (dynamic programming based) method to dissect any given protein structure using the inferred Proçodic dictionary.

These results are achieved due to the expressive power of tableaux to compactly represent the essence of protein folding patterns. This tableau-representation together with the statistically rigorous minimum message length inference methodology, provides a significantly more powerful framework to losslessly compress and identify redundancies in the protein folding space.

## Discussion

### Many concepts are linked to ligand-binding sites

The molecular function of proteins is often mediated via interactions with chemical components such as metal ions, coenzymes, metabolic substrates, and nucleic acids, amongst others. Knowledge of such interactions is central to annotate protein function (47, 48), engineer new proteins (49), and design novel drugs (50, 51). These functionally critical interactions impose structural constraints on protein structures, as their domains evolve from a common ancestor. As noted by Lesk and Chothia (52), in many cases active sites are the best-conserved regions within a family of protein structures (as seen in Fig. 4).

We have analysed our dictionary and systematically identified concepts directly related to protein-ligand interactions. To achieve this, we mined and catalogued frequent-ligand information (from ‘`HETATM`’ records) derived from the source `wwPDB` entries of each concept usage (i.e., each instance in the `wwPDB` where the concept is used in the dissection of that protein’s tableau). Our definition of a *ligand* comes from the inventory of 23,258 chemical components specified by the LigandExpo (53) database. We note that this inventory does *not* exclude simple monovalent ions (such as Na^+^, K^+^and Cl^*-*^) or those that are often not biologically functional (such as sulphate 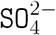 ions). To complement this information, we also mined and catalogued keywords (from ‘`KEYWDS`’ records) derived again from the source `wwPDB` entries of these concept usages. We used the observed frequencies of the bound ligands within the regions of concept usages, to narrow the initial set down to the 463 (31%) concepts that stand out in terms of recurrent patterns of interactions with the same set of ligand(s). These encompass interactions with monovalent ions, di-/tri-/tetra-valent ionic species, small molecules (including nucleotides), and macromolecular compounds, among others.

The full annotated list of concepts with observed interactions with ligands/chemical components is available in the supporting data file: `conceptsWithLigandInteractions.txt` (click).

Fig. 5 shows the distribution of 69 distinct chemical interactions observed within the shortlisted set of 463 concepts. Fig. 6(a-g) shows examples of concept usages for a random selection of 8 concepts associated with metal-binding activity. Table 1 shows a partial list of concepts for which all (100% of) their usages show binding to the specified ligand/chemical components. Also shown are the extracted high-frequency keywords associated with usages of that concept, providing useful insights to impute functional roles. Among the shortlisted set of 463 concepts are those that demonstrably show binding specificity linked with target recognition, reception and signalling (see Table 2).

**Table 1.**
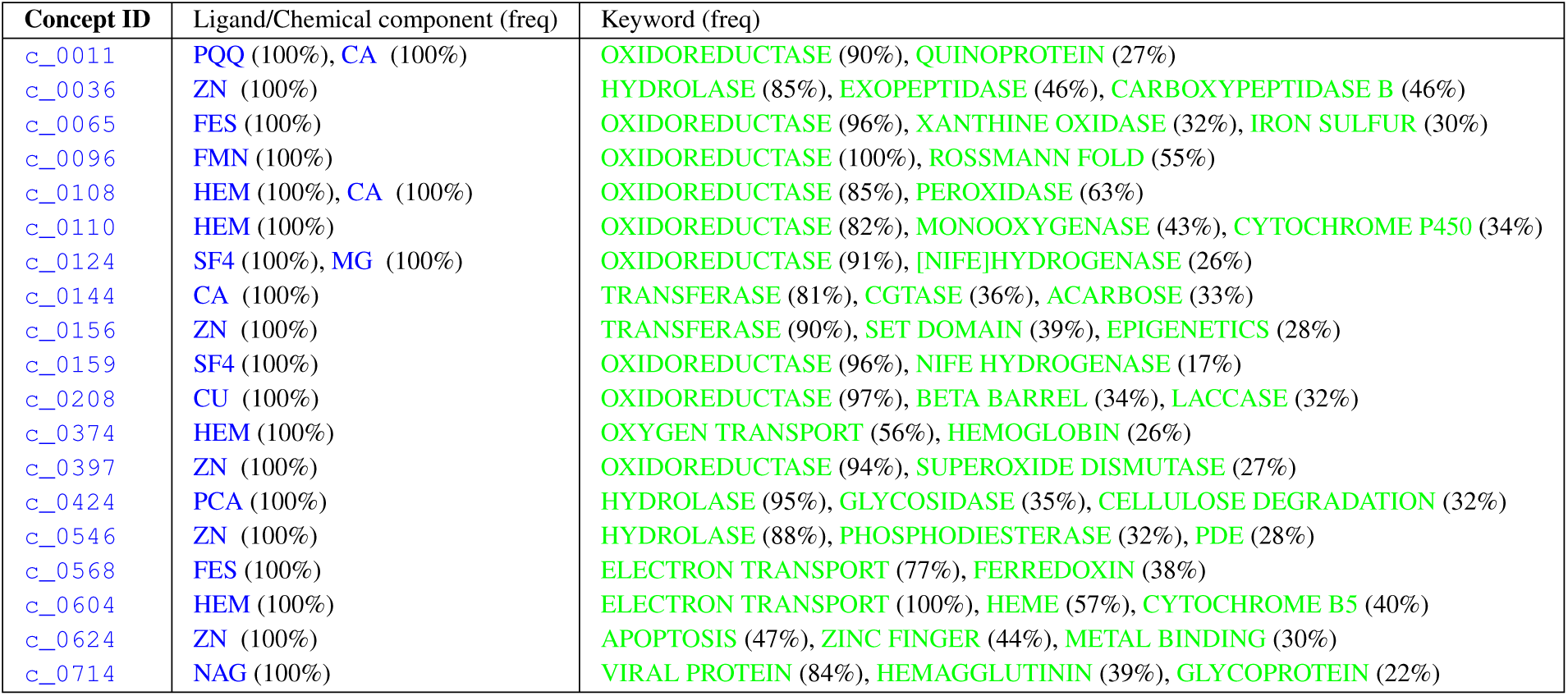
A partial list of concepts for which all (100% of) their usages show interactions with ligand/chemical components. This is derived by inspecting the ligand (‘HETATM’) records within the source coordinate files of each concept usage. The bound ligands are shown (in the second column) using their standardized abbreviations, along with their observed frequency within the usages in parentheses. Also shown (in the third column) are the top keyword terms (from ‘KEYWDS’ records specified by the structures’ authors) recurring within the usage coordinate files with their associated frequencies. (Note: CA = Calcium ion)

**Table 2.**
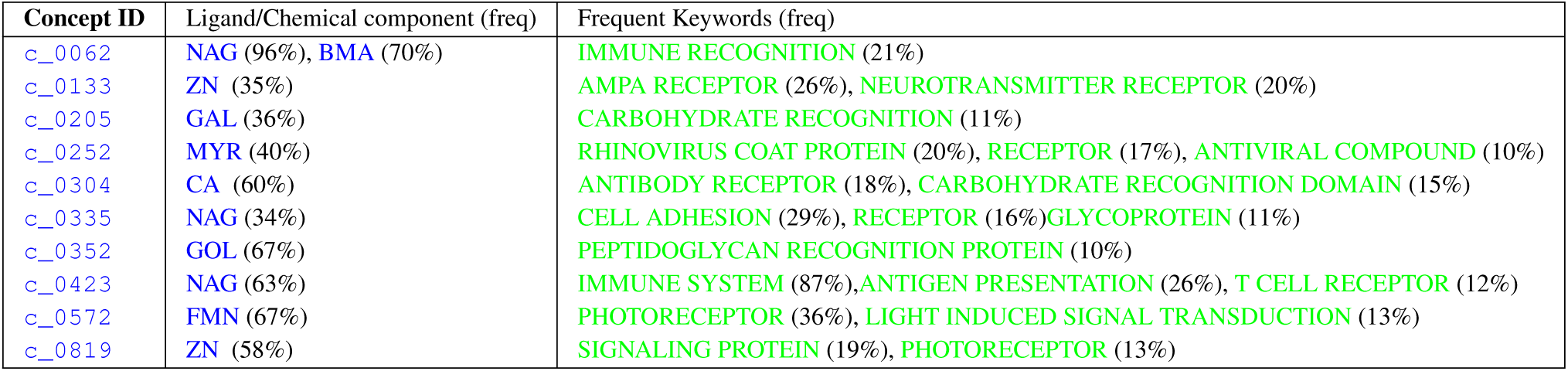
A partial list of concepts putatively linked to molecular reception, recognition, and signalling.

| Concept ID | Ligand/Chemical component (freq) | Frequent Keywords (freq) |
| --- | --- | --- |
| c_0062 | NAG (96%), BMA (70%) | IMMUNE RECOGNITION (21%) |
| c_0133 | ZN (35%) | AMPA RECEPTOR (26%), NEUROTRANSMITTER RECEPTOR (20%) |
| c_0205 | GAL (36%) | CARBOHYDRATE RECOGNITION (11%) |
| c_0252 | MYR (40%) | RHINOVIRUS COAT PROTEIN (20%), RECEPTOR (17%), ANTIVIRAL COMPOUND (10%) |
| c_0304 | CA (60%) | ANTIBODY RECEPTOR (18%), CARBOHYDRATE RECOGNITION DOMAIN (15%) |
| c_0335 | NAG (34%) | CELL ADHESION (29%), RECEPTOR (16%) GLYCOPROTEIN (11%) |
| c_0352 | GOL (67%) | PEPTIDOGLYCAN RECOGNITION PROTEIN (10%) |
| c_0423 | NAG (63%) | IMMUNE SYSTEM (87%), ANTIGEN PRESENTATION (26%), T CELL RECEPTOR (12%) |
| c_0572 | FMN (67%) | PHOTORECEPTOR (36%), LIGHT INDUCED SIGNAL TRANSDUCTION (13%) |
| c_0819 | ZN (58%) | SIGNALING PROTEIN (19%), PHOTORECEPTOR (13%) |

**Fig. 5.**
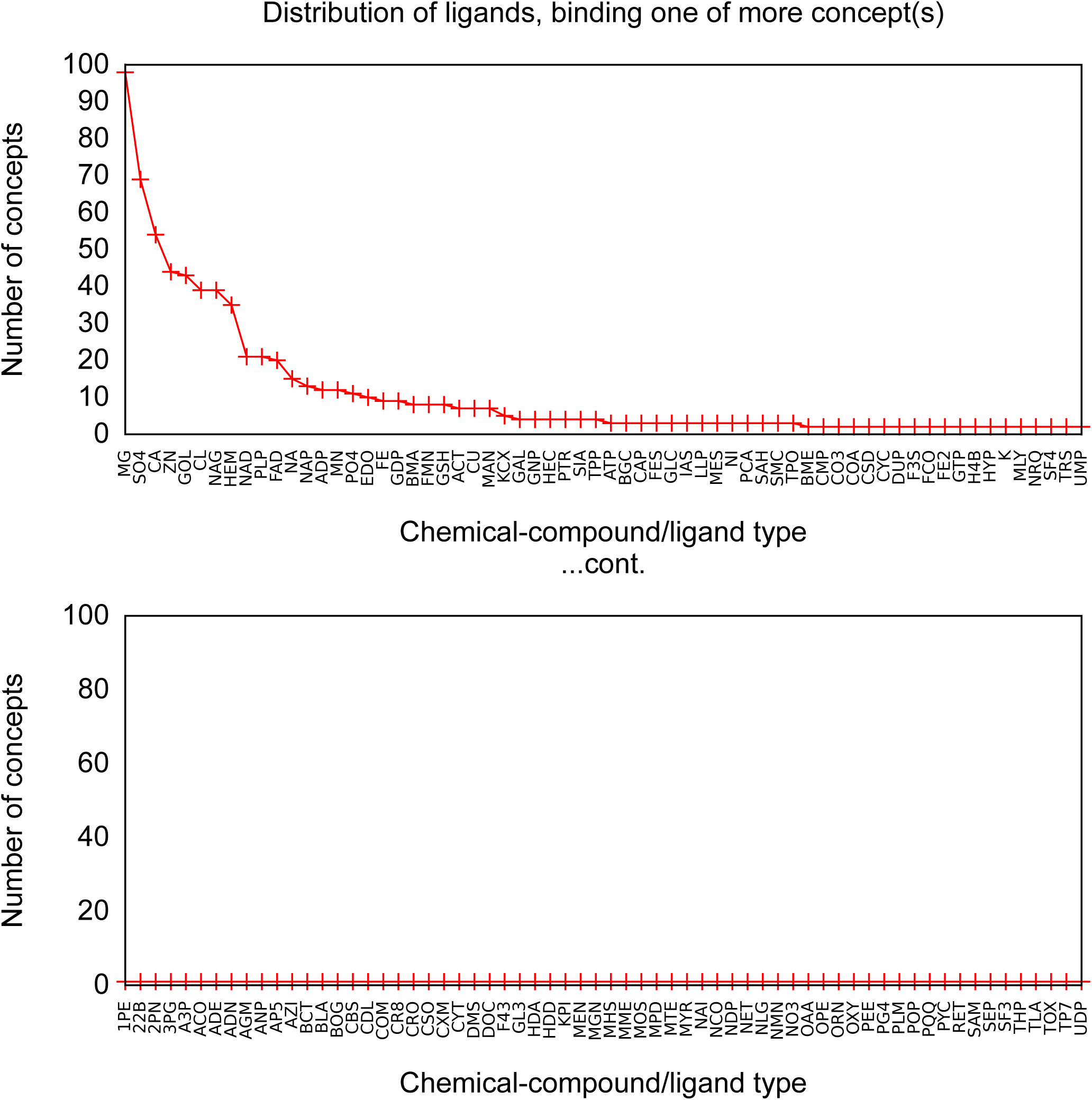
Distribution of 128 distinct ligands binding a shortlisted set of 463 concepts. A concept from the dictionary is shortlisted to be correlated to a binding site if > 30% of its usages in the `wwPDB` bind to a common ligand. The full details are available in `conceptsWithLigandInteractions.txt` (click). For readability, this distribution is split across 2 plots in the decreasing order of the number of shortlisted concepts that each ligand binds. In the top figure, the distribution varies between 98 (far left: `MG`) and 2 (far right: `UMP`). In the bottom figure, all ligands appear exactly in one of the shortlisted 463 concepts.

**Fig. 6.**
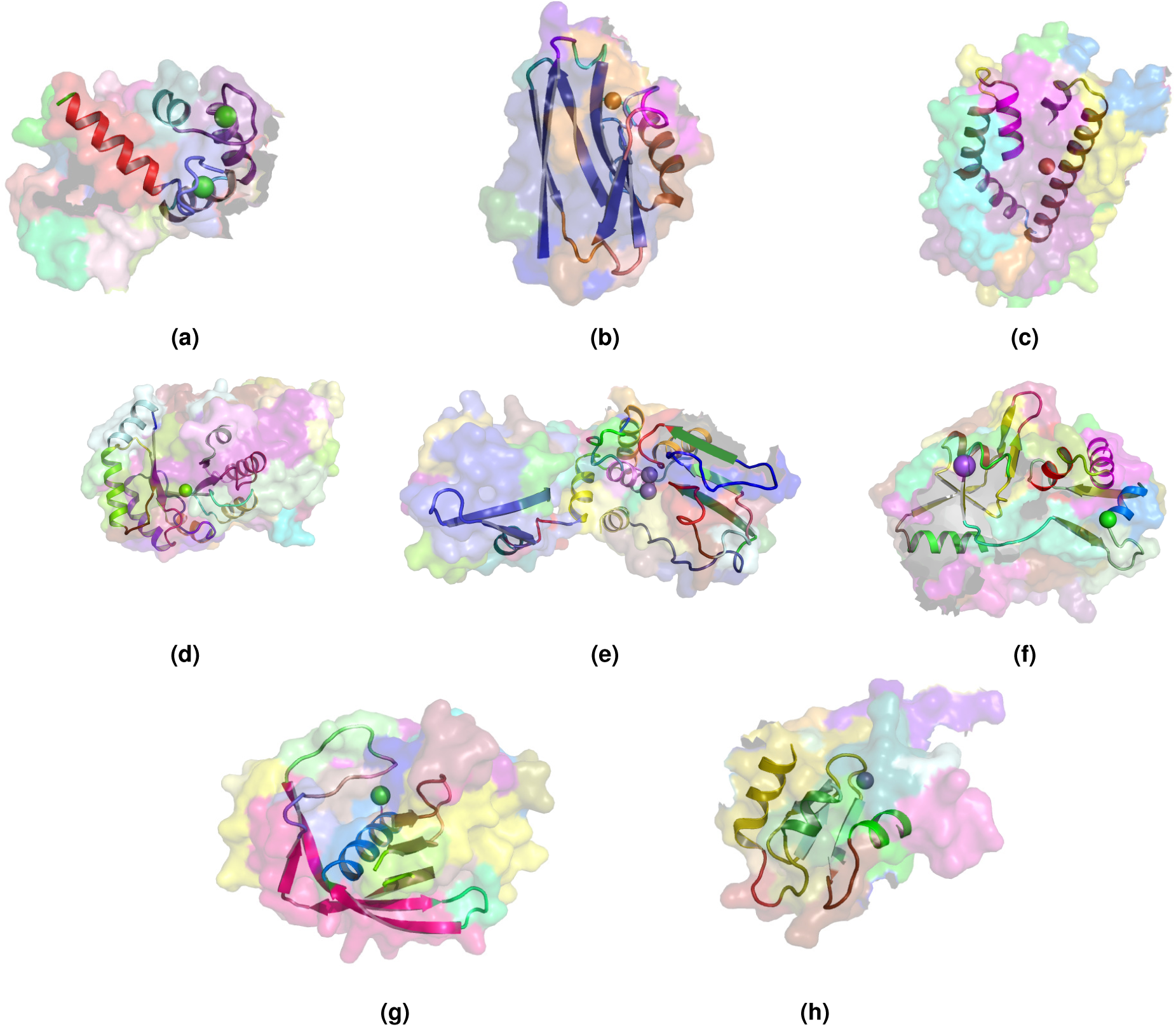
Exemplars of usages of eight concepts linked to metal binding activity. The region of concept usage is shown in cartoon in the context of the surface rendering of the source protein chain. (a) Usage of concept `c_1099` within the calcium-bound calmodulin (1CDL (54)). (b) Usage of concept `c_432` within the coppper-bound electron transfer protein (1A4B (55)). (c) Usage of concept `c_885` within the iron-bound oxidoreductase (2VUX). (d) Usage of concept `c_139` within the magnesium-bound lyase (3TTE). (e) Usage of concept `c_186` within the manganese-bound hydrolase (1K23 (56)). (f) Usage of concept `c_133` within the sodium cation-bound Kainate and AMPA receptors (3G3G (57)). (g) Usage of concept `c_280` within the nickel-bound peptide deformylase (2AIA). (h) Usage of concept `c_624` within the zinc-bound Melanoma inhibiting anti-apoptotic protein (1OY7 (58)).

The full list of inferred concepts putatively linked to molecular reception, recognition, and signalling is available in the supporting data file: `receptorConcepts.pdf` (click).

### Inferring biological function from concept usage information

Many proteins are deposited into the `wwPDB` with unspecified functional annotation, especially those coming from structural genomic initiatives. Functional characterization of such proteins is of crucial importance to the structural biology community. Its importance can be evidenced by the community-wide Critical Assessment of protein Function Annotation programme (CAFA, biofunctionprediction.org/cafa/), which assesses methods dedicated to predicting protein function from amino-acid sequence.

As previously shown (Fig. 4(a)), the rich source of information within this concept dictionary is useful to investigate and impute biological function. More evidence of this is shown by another case study involving the haze-forming thaumatin-like protein in white wines made from *Vitis vinifera* (4JRU containing 201 residues). Fig. 7 gives the dissection of 4JRU composed of two concepts `c_0111` and `c_1442`.

**Fig. 7.**
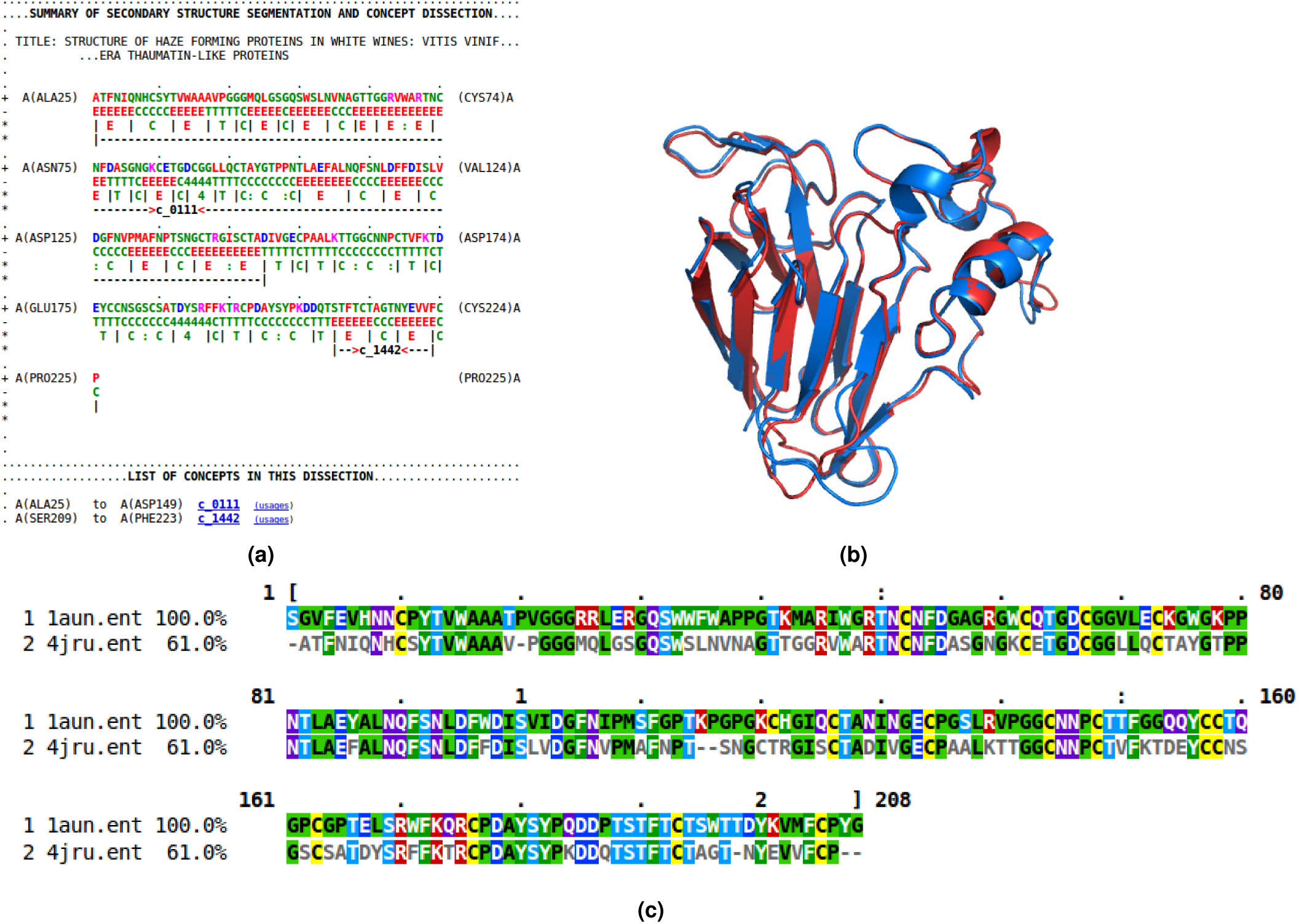
(a) Dissection output from Proçodic of the haze-forming thaumatin-like protein in white wines from *Vitis vinifera* (4JRU). (b) Superposition of this haze-forming protein and the pathogenesis-related PR-5d protein of tobacco (*Nicotiana tabacum*; 1AUN). 4JR is shown in blue; 1AUN is shown in red. This superposition was based on the structural alignment produced by `MMLigner` (59), which is shown in (c).

Concept `c_1442` is of less functional interest as it defines a common *β*-hairpin unit consisting of two antiparallel *β*-strands. On the other hand, `c_0111` contains 12 strands that assemble to form mainly two face-to-face packed antiparallel *β*-sheets with an extended *β*-ribbon connected by an Ω-loop (60). This multi-stranded motif is characteristic of thaumatin-like proteins (61). Examining the usages of this concept within the `wwPDB` via our Proçodic web site, we find it is used at 15 other loci, most of them thaumatin/osmatin-like proteins, with their top two keywords displaying ‘`antifungal protein` (53.3%)’ and ‘`plant protein` (46.7%)’, respectively. Fig. 7(b-c) show the structural alignment of 4JRU with the usage in the pathogenesis-related PR-5D protein of tobacco (*Nicotiana tabacum*; 1AUN with 208 residues) that results in a superposition with 1.47 Å root-mean-square deviation (r.m.s.d.) over 201 amino acid residues between the C_*α*_ coordinates of the two structures. This specific PR-5D protein is classified functionally as an antifungal protein and, in general, proteins of this class have known pathogenesis-related antifungal activity. This suggests that the haze-forming protein might exhibit the same biological function.

In some cases, the information provided by this dictionary can lead to a reliable but less-specific functional classification prediction, for example putatively identifying a general type of function such as ‘Oxidoreductase’ or ‘Lyase’. Such generic functional classification can be useful, as it may provide guidance for laboratory experiments aimed at defining the function more precisely, especially if clues about a ligand-binding site are available. For example, consider the crystal structure of dihydrodipicolinate synthase (DapA) from *Agrobacterium tumefaciens* (2HMC). The dissection of this DapA structure shows the usage of concept `c_0008` covering its entire chain A. About 90% of `c_0008`’s 118 usages show the functional classification as ‘Lyase’. DapA belong to the family of amine-lyases that catalyze the cleaving of carbon-nitrogen bonds, playing an important role in lysine biosynthesis in prokaryotes, phycomycetes, and plants (62). A similar example would be the identification of HI0073 from the *Haemophilus influenzae* structural genomics project as a nucleotide-binding protein (63).

### Local sequence-structure correlation within concept usages

The identification of structural features that have strong amino-acid sequence preferences is central to structure prediction (45). Therefore, we studied the concept usages within the `wwPDB` to explore the conformational preferences of local sequences. To achieve this, for each concept, the amino acid sequences in the regions of concept usages within the `wwPDB` were extracted, and the sequences in each set were aligned and clustered (64).

Almost 20% of the concepts in our dictionary (288 out of 1,493) have associated amino acid sequences that cluster into a single group. Fig. 8(a) shows the number of clusters produced for each set of concept usage sequences – in general, the fewer the clusters, the stronger the local sequence-structure relationship. When considering the (normalized) ratio of clusters over the number of *non-identical* amino acid sequences of concept usages (Fig. 8(b)), almost 30% of the concepts (441) have a ratio smaller than 0.05, while almost 50% (738) have a ratio between 0.05 and 0.1. Together, this indicates that for almost 80% of the concepts in our dictionary, their usages of amino-acid sequences cluster into a small number of groups (*<* 10% of their total unique amino acid sequences).

**Fig. 8.**
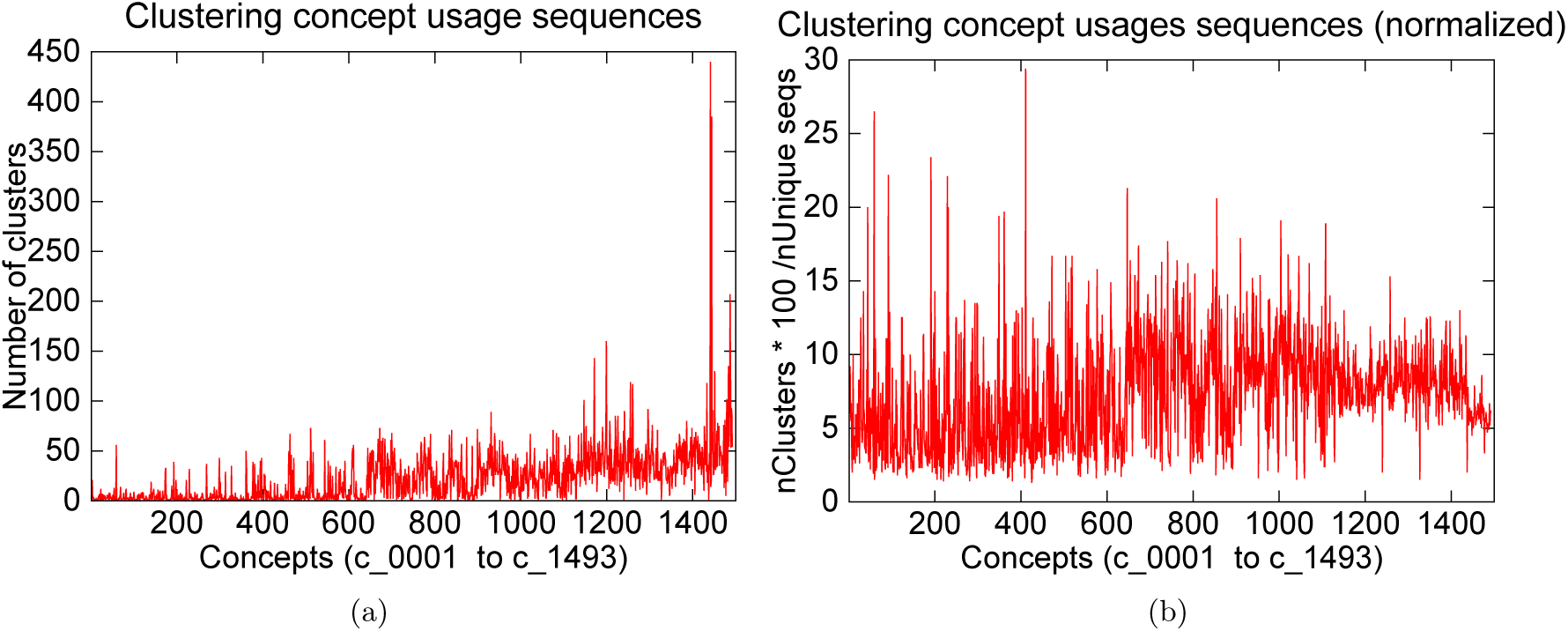
(a) Number of clusters produced by `Clustal-Omega` (64) based on its computation of the multiple sequence alignment of the usage-amino-acids-sequences for each concept. `Clustal-Omega` uses the `mBed` algorithm to cluster sequences (65). (b) Normalised plot to account for the differences in the number of usages per concept. Normalization involves dividing the number of clusters by the total number of *unique* amino acid sequences observed in the set of usages per concept.

This strong sequence dependence is expected, particularly for concepts linked to ligand binding or other functional units. For example, Fig. 9 shows the sequence logo obtained from the multiple sequence alignment of the usage sequences of the concept `c_0397`. This concept is related to the Cu-Zn type I (SODI) superoxide dismutase, which has a *β -* barrel-like subunit with copper and zinc ions bound at the active site. This is common in many Gram-negative bacterial pathogens (amongst others) to counteract a burst of toxic superoxide radicals under oxidative stress (66).

**Fig. 9.**
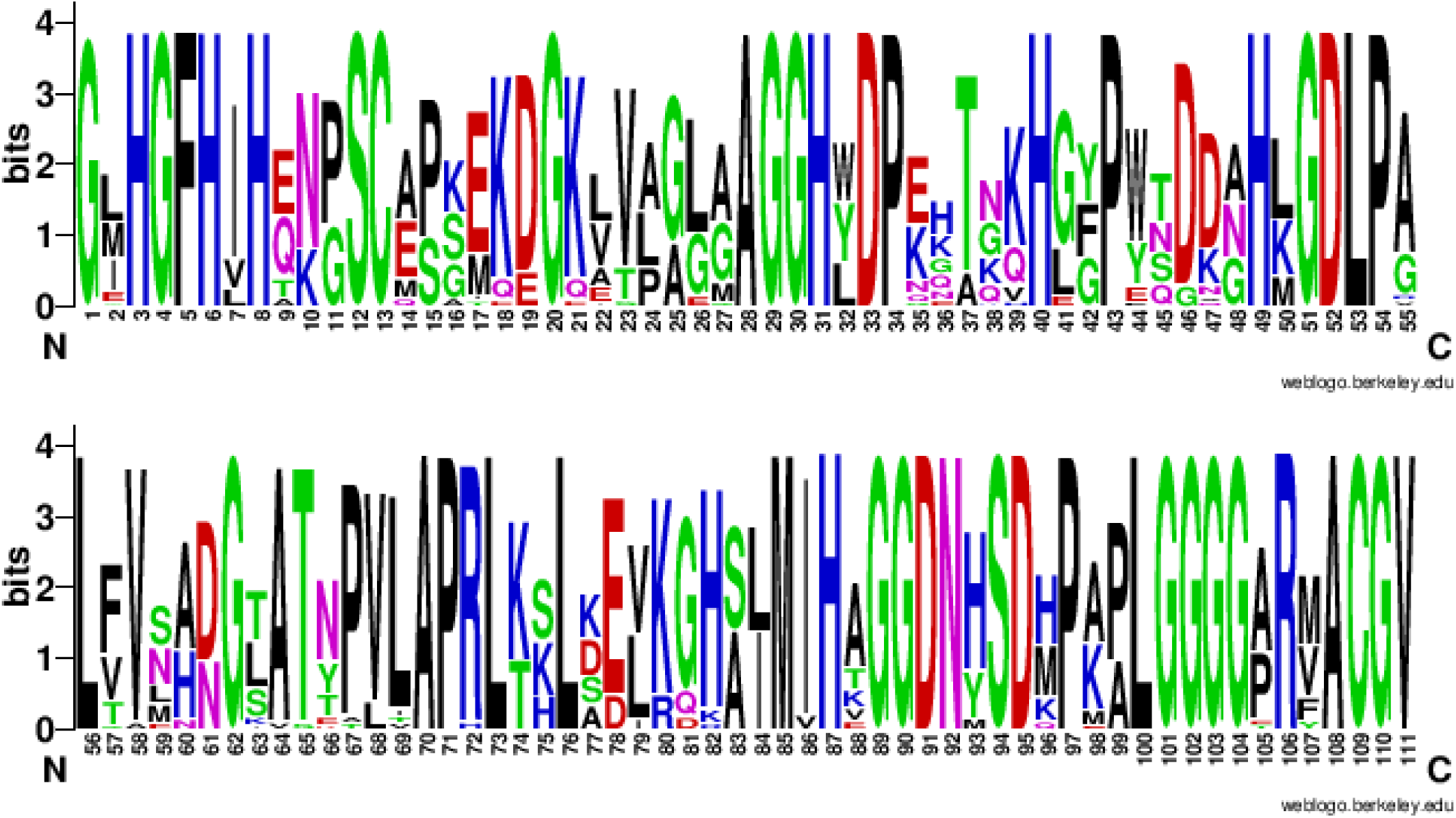
Amino acid sequence logo (in two parts: columns 1-55 and 56-111) showing the sequence consensus across the usages of a randomly chosen concept `c_0397` directly related to the Cu-Zn binding superoxide dismutase. Of the 111 columns in the multiple sequence alignment (of the `c_0397`’s 33 usage sequences) corresponding to this logo, 46 aligned columns show a consensus of 100%.

There is a potential application of the observed sequence-structure correlations to structure prediction. We downloaded the coordinate files of 33 `wwPDB` structures specified in the description field of the CASP12 target list available at http://predictioncenter.org/casp12/targetlist.cgi. Each chain from these 33 structures was independently dissected using the Proçodic dictionary of concepts. The dissection of protein chains defines non-overlapping regions assigned either to one of the dictionary concepts (`c_0001` – `c_1493`), or a *null* concept (`c_0000`). For each region assigned to a dictionary concept, we extracted the associated target amino-acid sequence and performed a pairwise *sequence* alignment with each of the local amino acid sequences defined by the concept usages. This exercise identified a subset of concept usages in the `wwPDB` whose local amino acid sequences have a detectable similarity with the target. Table 3 quantifies the extent of coverage of these regions for each of the 33 CASP12 targets. This table shows that in 26 of 33 cases, more than 50% of the target amino-acid chain has detectable sequence similarity that can be derived from the usage information.

**Table 3.** Statistics showing the extent of detectable sequence similarity on each of the 33 CASP12 targets with their wwPDBIDs specified at http://predictioncenter.org/casp12/targetlist.cgi. First column: wwPDBID of the 3D experimental structure of each CASP12 target. Second column: The coverage statistics in terms of the total number of amino acids (#a.a.) within the amino acid (sub-)sequences defined by the dissected regions of the target protein with detectable sequence similarity with amino acid (sub-)sequences of their corresponding concept usage instances (see main text). Third column: The total number of amino acids in the target protein, cumulative over all chains. Fourth column: Percentage coverage = Second column*100/Third column.

| Target's wwPDBID | #a.a.'s in regions with usage seq hits | Total #a.a.'s in all chains | %Covered |
| --- | --- | --- | --- |
| 3JB5 | 1046 | 2076 | 50.4 |
| 4YMP | 202 | 215 | 94.0 |
| 5A7D | 4468 | 5065 | 88.2 |
| 5AOT | 91 | 102 | 89.2 |
| 5AOZ | 125 | 141 | 88.7 |
| 5D9G | 154 | 502 | 30.7 |
| 5ERE | 417 | 540 | 77.2 |
| 5FHY | 155 | 458 | 33.8 |
| 5FJL | 88 | 136 | 64.7 |
| 5G3Q | 100 | 168 | 59.5 |
| 5G5N | 580 | 1022 | 56.8 |
| 5HKQ | 160 | 263 | 60.8 |
| 5IDJ | 63 | 242 | 26.0 |
| 5J4A | 339 | 440 | 77.0 |
| 5J5V | 815 | 1065 | 76.5 |
| 5JMB | 103 | 182 | 56.6 |
| 5JMU | 172 | 219 | 78.5 |
| 5JO9 | 215 | 239 | 90.0 |
| 5JZR | 203 | 262 | 77.5 |
| 5KKP | 166 | 509 | 32.6 |
| 5KO9 | 73 | 253 | 28.9 |
| 5LEV | 323 | 375 | 86.1 |
| 5M2O | 171 | 211 | 81.0 |
| 5MQP | 2674 | 4801 | 55.7 |
| 5NSJ | 150 | 284 | 52.8 |
| 5NV4 | 713 | 1377 | 51.8 |
| 5SY1 | 786 | 1458 | 53.9 |
| 5T87 | 444 | 745 | 59.6 |
| 5TF2 | 331 | 338 | 97.9 |
| 5TJ4 | 2640 | 5462 | 48.3 |
| 5UNB | 378 | 681 | 55.5 |
| 5UVN | 954 | 2496 | 38.2 |
| 5UW2 | 211 | 332 | 63.6 |

It should be noted that we used the structural information of CASP12 targets to dissect the protein chains, before identifying the sequence relationships of the target sequence and those within the concept usages. However, for the proper application to structure prediction, the identification of sequence hits with concept usages should be carried out using only the target sequence. In principle, this can be done by sliding along the target sequence with varying window sizes, and exploring the sequence similarity with the sequences across all usages of every concept in the dictionary. Nevertheless, this preliminary analysis can be used to hypothesise reasonably that these local sequence-structure relationships provide a strong potential to support structure prediction efforts, especially since an average concept usage spans significantly longer stretches along the protein chain than the currently considered oligopeptide fragment libraries used by fragment-based *ab initio* protein modeling approaches. Thus, this information can be potentially utilized to model several non-overlapping regions in the target protein chains by the current state-of-the-art structure prediction servers (67–69).

The amino acid subsequences of non-overlapping regions dissected using the Proçodic dictionary of concepts is available at: `casp12_prosodic_dissections.tgz` (click). The information of dissected target region followed by other subsequences in the usages of the corresponding (assigned) concept with demonstrable sequence similarity (under pairwise sequence alignment with the target subsequence) is available at: `casp12_concept_usage_hits.tgz` (click). The multiple sequence alignments (using MUSCLE (70) with default options) of the identified sequence hits is available at: `casp12_concept_usage_hits_msa.tgz` (click).

### Exploration of substructures and structural relationships

In addition to the applications explored above, the dictionary can be used to complement standard protein structural studies. Researchers can approach the dictionary with a particular structure or family of structures in mind. For example, dissecting the human haemoglobin (1HHO, chain A) at the Proçodic web site identifies the concepts `c_0375`, `c_0894` and `c_1410`. Choosing one of the concepts, for example `c_0894`, its archetype is found in `d1×9fd`, a globin from the annelid *Lumbricus terrestris*. Note that related proteins can present dissections into different concepts. However, these concepts can still be related (see discussion on hierarchical clustering of concepts on Page 5). For example, `c_0375` and `c_0894` are related concepts linked to globins, with the former being more elaborate (with three extra helices) than the latter. Examining the corresponding concept ‘usages’ link on the Proçodic web site reveals that many usages of these related concepts appear in other globins. **Supplementary §S3** contains several examples of use of Proçodic to explore protein substructural similarities.

## CONCLUSION

Most protein domains fold into geometrical assemblies of helices and strands of sheet. Our work has analysed the domains of known structures and inferred a ‘basis set’ containing 1,493 folding concepts into which folding patterns of domains can be dissected. The discovery of the dictionary was completely automatic, and unbiased by any previous structural analysis of folding patterns, or by any hidden sequence or structure-based patterns. The effectiveness of our inference method is validated by the discovered concepts, which subsume classic supersecondary structures (*α*-hairpin, *β*-hairpin, and *β*-*α*-*β* unit etc.), known repeat patterns (ankyrin, armadillo, kelch repeat etc.) and many other known patterns (*β*-propeller, Jellyroll, and Immunoglobulin architecture, among others). We note that the scope of the concepts in our dictionary far exceeds these known patterns.

The discovery of this dictionary allows us to dissect optimally the structures of any protein domain in seconds, and map dictionary concepts onto non-overlapping regions along its chain. Importantly, the dissections of domains into concepts provide a plethora of useful biological insights, including:

1. Understanding the fundamental components of protein folding patterns. Our dictionary of concepts will support innovative projects aimed at the analysis of protein structures.
2. Correlation, in many cases, of concepts with functions directly or indirectly, via ligand binding sites. This provides useful predictions in the case of proteins with known structure but unknown function.
3. Many concepts show amino-acid sequence correlation; that is, some conservation of sequence patterns. These results are applicable to protein structure prediction by suggesting conformations of local regions.

The results of dissecting all structures in the current `wwPDB`, or of dissecting a user-supplied set of protein coordinates, are accessible from the Proçodic web site: http://lcb.infotech.monash.edu.au/prosodic (click). This site supports the interactive exploration of protein structures and their relationships.

## SUPPLEMENTARY MATERIAL

Supplementary information, including description of material and methods supporting this work, is available as a separate PDF.

## ACKNOWLEDGEMENTS

This research is funded by Australian Research Council (ARC) Discovery Project grant (DP150100894). We thank Research Computing Centre, University of Queensland for the High-Performance Cluster Infrastructure that supported this project over the last 3 years. AML thanks the Medical Research Council Laboratory of Molecular Biology for their hospitality during his sabbatical year. We thank Sureshkumar Balasubramanian for proofreading this work.

## S1. Material and Terminology

### Tableau representation of protein folding patterns

As mentioned in the main text, the essence of any protein folding pattern can be captured by the order, geometry and contact information of its secondary structural elements (1). Lesk (2) developed the *tableau* representation to capture this information in the form of a concise symmetric matrix (see Fig. 1 in main text). A tableau encapsulates: (1) the order in which helices and strands-of-sheet appear in the protein chain, represented by a string, **S**, of length |**S**| over the {H(for helix), E(for strand)} alphabet; (2) the geometry of each pair of secondary structural elements, represented by a square-symmetric matrix of angles, **Ω**, of order |**S**| × |**S**|, where angles are in the range (*-*180*°,* 180*°*]; (3) the corresponding interactions between pairs of secondary structural elements, represented by a contact matrix, **Ξ**, of 0/1 values and order |**S**| × |**S**|, where 1 represents contact and 0 otherwise (3, 4). Formally, any tableau *τ* is a three-tuple of the form (**S**, **Ω**, **Ξ**).

### Source collection used for the inference of PROçODIC dictionary of concepts

A *source collection* is a collection of (source) tableaux, denoted by the set 𝒯= {*τ*_1_, *τ*_2_, …, *τ*_|𝒯|_*}*.

Since the full `wwPDB` has a lot of redundancy in terms of entries with similar structures, to infer the dictionary of concepts, we use the ASTRAL SCOP-95 (5–7) (v2.05) dataset which has been produced to remove bias due to over-represented structures, while explicitly incorporating structure quality at each step of the domain selection. (8). This data set used is composed of 26,949 domains, representing only 12% of the full SCOPe (v2.05) domain dataset. Of these, 13,365 domains have < 40% sequence similarity. Although the maximum sequence similarity two proteins can share is 95%, the average sequence similarity is significantly lower (< 53%). The full list of ASTRAL SCOP-95 domains used to infer the reported dictionary is available in the supporting data file: `prosodicInferenceList.txt` (click).

### Subtableaux and the dictionary of concepts

PROçODIC concepts are represented as contiguous subtableau. A candidate concept, denoted by *c*, can be instantiated by selecting a source tableau *τ*_*𝓁*_ *∈*𝒯 and specifying a continuous range of indices [*i, i* + 1, …, *j -*1, *j*], such that 1 ≤*i* ≤ *j ≤*|*τ*_*𝓁*_ |, provided it satisfies the constraint that the graph defined by the corresponding contact matrix is connected – any undirected graph is said to be connected if there exists a path between any two vertices in the graph.

Formally, any concept 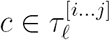 defines a three-tuple 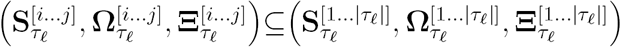 Here, 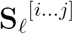 is the substring of **S**_*𝓁*_, defined by the range [*i,* …, *j*], while 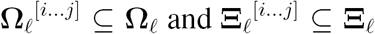 are the corresponding geometry and contact submatrices respectively. We define a *dictionary* as a set of concepts, denoted by the set *𝒞* = {*c*_1_, *c*_2_, …, *c*_|*𝒞*|_}. A dictionary can contain an arbitrary number of concepts (|*𝒞*|), with each concept *c*_*y*_ *∈𝒞* containing an arbitrary number of secondary structural elements (|*c*_*y*_| ≥ 2). Any possible dictionary is a potential candidate to compress the source collection of tableaux, 𝒯.

Associated with each concept *c*_*y*_ *∈𝒞* is a concentration parameter, *κ*_*y*_, corresponding to a von Mises circular (angular) probability distribution (9). This parameter controls the assignment of probabilities used to estimate the encoding length of entries in **Ω** when compressing regions of the source tableaux. That is, *κ*_*y*_ controls the *flexibility* of an inferred concept. A smaller/larger *κ*_*y*_ yields greater/lesser flexibility of the concept’s usages for compressing source tableaux regions. In this work, all {*κ*_*y*_*}*_*∀*1 *≤y ≤*|*𝒞*|_ lie in the range [*κ*_min_ = 10, *κ*_max_ = 100], with their precise value inferred (to a precision of *ϵ*_*κ*_ = 0.5) as a part of the dictionary search (see below).

### Collection of wwPDB files dissected using this dictionary

At the time of writing this article, PROçODIC includes, collectively within the usages of concepts, the dissections of 113,724 protein coordinates files from the `wwPDB` (10). The specific list is available in the supporting data file: `prosodicDissectedWWPDBList.txt` (click). In addition to these dissections, the PROçODIC website allows users interactively to dissect any protein structure on demand.

## S2. Methods

### Secondary structure assignment and construction of tableaux

Secondary structure is assigned to any given protein coordinate data using our algorithm, SST (11). SST assigns secondary structures using solely the C_*α*_ coordinates of protein structures. The web version of SST is available from: http://lcb.infotech.monash.edu.au/sst. Using SST we can delineate any protein structure into its standard secondary structural elements (SSEs) and thence generate its tableau representation.

### Inference of dictionary of concepts

In the proceedings of 2017 IEEE Data Compression Conference, we described the compression-based methodology of inference of recurrent subtableaux on any source collection of tableaux using the Bayesian method of Minimum Message Length (MML) inference (12). The dictionary we report and analyse in this work has been inferred using this methodology with minor modifications. For convenience of the reader, the description of our methodology is reproduced in the Appendix (see Page 9).

## S3. Supplementary Notes for ‘Exploration of substructures and structural relationships’

### Globins

As seen in the main text, different related proteins can present dissections into different related concepts; this is the result of the calculation to optimize the representation of the whole set of proteins. For closely-related proteins, the three-dimensional structures of the usage instances of a given concept are superposable. Fig. SF1(a) shows the superposition of the instances of `c_0894` from the *α* subunits of human deoxyhaemoglobin (2DN2), and oxyhaemoglobin from the common pigeon *(Columba livia)* (2R80). In the structural superposition, 78 C*α* atoms from these regions fit to an r.m.s.d. of 0.83 Å.

**Fig. SF1.**
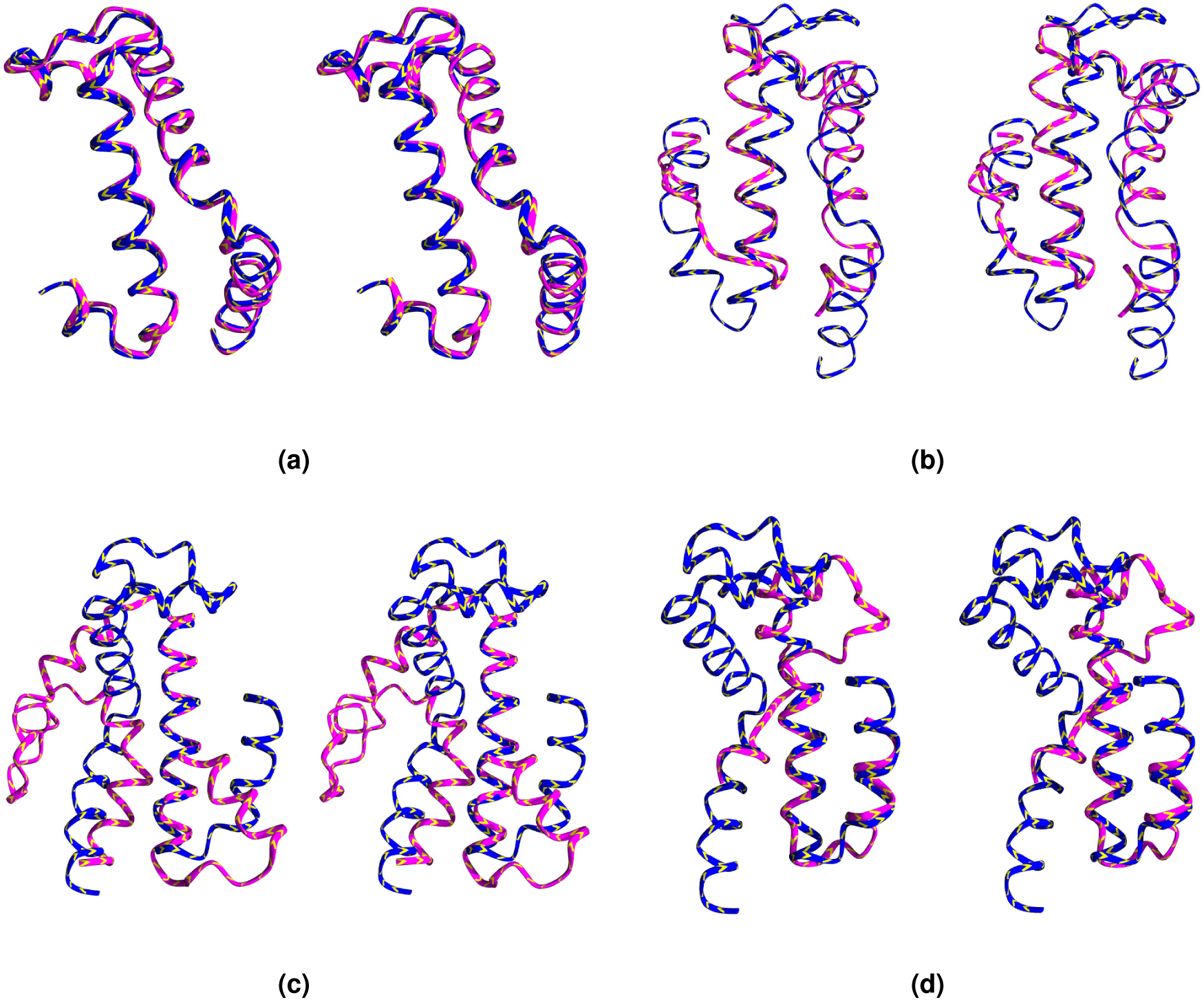
Superposition of the usage instances of concept `c_0894` (shown in stereo). (a) Superposition of instances from the *α* chains of human deoxyhaemoglobin (blue) and oxyhaemoglobin from common pigeon *(Columba livia)* 2R80 (pink). (b) Superposition of instances from human oxyhaemoglobin (1hho) and the truncated globin of *Tetrahymena pyriformis* 3AQ5. (c) Superposition of instances from human haemoglobin (1HHO chain A) with the corresponding usages in an *unrelated* protein, complex II (succinate dehydrogenase) from *E*. *coli* (1NEN chain B). (d) Superposition of 1HHO with unrelated human squalene synthase (3VJ8).

More distantly-related globins may also share the concept but the regions are not so precisely superposable. A structurally-diverged class of globins are the truncated globins, substantially shorter than sperm-whale myoglobin and showing substantial structural changes. Fig. SF1(b) shows the superposition of the instances of `c_0894` from human oxyhaemoglobin (1HHO) and the truncated globin from the ciliate *Tetrahymena pyriformis* (3AQ5). Note that the helix lengths are much more variable than in the superposition shown in Fig. SF1(a). The loop region at the top of this figure does not superpose well between the two structures, and indeed does not even have the same length. This emphasises that our representation of folding patterns captured via the subtableaux of concepts is at the level of geometry of secondary structure elements.

The list of `wwPDB` entries reported from the ‘usage’ link contains proteins identified as non-globins, for instance complex II (succinate dehydrogenase) from *Escherichia coli,* a membrane protein, (1NEN chain B residues ASP144–LEU197). Superposing the residues in this region with the instance in 1HHO gives the result shown in Fig. SF1(c). The helices from the N and C termini of these regions fit well. The intervening region shares the secondary structure with some conformational differences.

Could it be that *Escherichia coli* complex II (succinate dehydrogenase) is really homologous to the globins? Examination of 1NEN shows that this protein includes strands of *β*-sheet, ruling out the possibility of similar topologies of their overall folding patterns.

Another example that shares a concept with human haemoglobin, is the human squalene synthase (3VJ8). (Fig. SF1(c)) Comparing the superpositions within the globin family with the superpositions involving globins and non-globins, closely-related globins show a well-fitting superposition of all secondary structures using the same overall rotation and translation, but globin–non-globin superpositions do not. This is because preserving the angles between successive secondary structures does not fix the global structure, although it does constrain it.

### TIM-barrels

When probing the web site for a standard type of structure, the TIM barrel, we enter into the keyword window the string EHEHEHEHEHEHEHEH – or its regular expression: `(EH){8}` – signifying an eight-fold repeat of a *β*-*α* unit.

The web site returned two concepts, `c_0008` and `c_0032` containing the pattern EHEHEHEHEHEHEHEH. The first, `c_0008`, contains TIM barrels. The structure of the concept comprises the canonical 8-fold *β*-*α* barrel plus four additional C-terminal helices. As the reader is encouraged to try, clicking on the image produces a large ‘still’ high-quality graphic display; clicking on ‘view interactively’ produces an image rotatable under mouse control. Clicking for ‘full details’ gives the full secondary-structure assignment for the ‘fold archetype’ of `c_0008`, SCOP domain `d3flua_`, and the tableau computed for this domain. (Our methodology enables each concept in the dictionary to converge to an archetype that can viewed as the topological *median* over all usages of that concepts in the source collection the dictionary compresses. Varying ideas defining such topological medians representing supersecondary structural motifs were previously explored, especially as ‘attractors in fold space’ to enable protein structural classification efforts (13, 14).

Clicking on ‘usages’ reports other instances of this concept in the `wwPDB`. There are 118 other usages, all TIM barrels. However, they are not the only TIM barrels in the `wwPDB`. Many structures that do contain the 8-fold barrel but lack the four C-terminal helices are dissected into smaller units, some containing *β*-*α* subsets of the barrel. Others proteins have helices inserted at different points, deviating from the specified secondary structural pattern. Therefore, searching using patterns alone does not return all the TIM barrels in the `wwPDB`.

It is possible to type ‘TIM barrel’ into the keyword field. The web site will then return many concepts, some of which do correspond to TIM barrels, but others of which do not. For instance, in response to the keyword query ‘TIM barrel’ Proçodic returns `c_0004` which contains 24 consecutive *β*-strands but no helices. The reason is that one of the `wwPDB` entries in which `c_0004` appears is the human PRMT5:MEP50 complex (4QGB): this entry protein does contain a TIM-barrel domain (in which `c_0004` does not appear), and the entry file contains TIM BARREL in its `wwPDB` KEYWDS record line, which triggers a hit on this concept. In summary, to find TIM barrels in the `wwPDB`, a Proçodic search for (EH){8} returns too little, a search for ‘TIM BARREL’ returns too much, and there is no Goldilocks compromise. In any event, this is a solved problem. For this particular question, other tools such as SCOP are more convenient and appropriate.

The other concept returned for the query EHEHEHEHEHEHEHEHEH, contains two domains, each with four *β*-*α* units, but not closed into a barrel. This appears because a sequence of secondary structure elements does not uniquely define three-dimensional structure.

### Uncompressed regions in dissections and unusual structural components

Suppose a region in a protein contains an unusual conformation. It may require a shorter message to send the subtableau information of this region raw (without compressing that region) than to include a representative within the dictionary and compress it. This is because, given its rarity, the overhead of adding it in the dictionary does not justify its inclusion as a separate concept. Overall, in the dissections of *∼*114 000 `wwPDB` entries, *∼*66% of the residues are covered by dictionary concepts.

An example of an uncompressed region appears in the dissection of *Chironomus* erythrocruorin, a globin from an insect. Although the overall structure of this molecule is similar to that of globins, there are deviations from the usual structure in the region corresponding to the D helix of sperm whale myoglobin. This results in different dissections of sperm whale myoglobin (1MBD) and *Chironomus* erythrocruorin (1ECD):

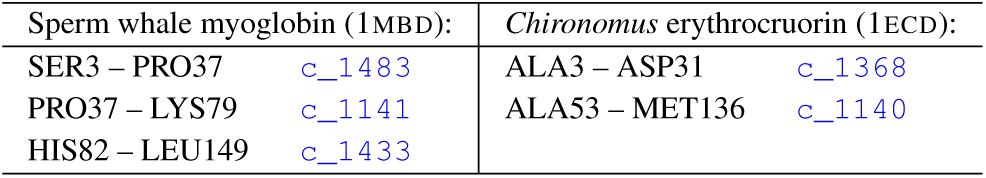

Observe that the region in *Chironomus* erythrocruorin from residues 31–53 is not part of the dissection. This region is explained directly without compression, using what we call the *null* concept.

### Erythrocruorins

Typing ‘erythrocruorin’ in the keyword field returns 26 concepts. The top one, concept `c_0547`, is an assembly of seven helices, corresponding to helices A-B-C-E-F-G-H in the canonical globin fold. There are 268 usage instances of this concept, which include erythrocuorins and globins. (The seven helix pattern fits the *α -*chain of mammalian haemoglobins, which lack a D helix, but not the *β*-chain which contains a D helix.) The distinction between similar structures that differ in some detail which breaks a pattern, can be seen as both a strength and as a weakness. We saw a similar phenomenon with the TIM barrels and with globins.

Substructures of the ‘globin fold’ containing 6 helices, include `c_0640`. This corresponds to globin helices A-B-E-F-G-H. Examples include phycocyanins and phycoerythrins, colicin, and certain globins (15). To be a proper instance of `c_0640`, a globin must *lack* both C and D helices. In these cases the region corresponding to the D helix is not helical, but the region corresponding to the C-helix is nevertheless quite close to the expected 3_10_ helix. This is because the region is distorted enough to drag at least one hydrogen bond outside the thresholds of acceptance in distance and/or angle. As a result, the region is assigned as a coil, and only six helices are attributed to the chain. In other cases, the compression criterion encodes a regular globin structure as more than a single concept. Thus, human oxyhaemoglobin (1HHO), chain A, is decomposed into:

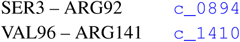

Note that this chain does have a 3_10_ C helix, but *not* a D helix.

In addition to helical concepts that are substructures of the canonical globin fold, querying for erythrocruorins (as a keyword on the web site) returns concepts containing purely *β*-sheet concepts. These are known to appear in large, multimeric, extracellular invertebrate erythrocruorins, as linker regions between globin-like all-helical domains (16). Our dictionary has therefore called for attention to these additional substructures, not customary structure components from the familiar globin family.

As a *β*-sheet structure is rare in the globins and erythrocruorins, it is of interest to explore the relationships of these substructures to other families. Are there homologues, of which the erythrocruorin linker domain might be part of a chain of evolutionary relationships?

Consider the concept `c_0559`, comprising six *β*-strands. Its *usages* provides a list of chains in which this concept occurs. Checking the list of chains against the SCOP classification gives the following results:

| #Instances | SCOP classification | #Instances | SCOP classification |
| --- | --- | --- | --- |
| 39 | b.60.1.2 | 1 | d.129.3.8 |
| 5 | e.7.1.1 | 1 | d.129.3.5 |
| 3 | h.1.2.1 | 1 | b.97.1.1 |
| 3 | b.8.1.1 | 1 | b.82.1.23 |
| 3 | b.30.5.4 | 1 | b.82.1.11 |
| 2 | d.85.1.1 | 1 | b.61.7.1 |
| 2 | d.25.1.1 | 1 | b.61.1.1 |
| 2 | b.60.1.1 | 1 | b.60.1.8 |
| 1 | h.1.32.1 | 1 | b.60.1.7 |
| 1 | g.12.1.1 | 1 | b.23.1.1 |
| 1 | f.1.1.1 | 1 | b.1.9.2 |
| 1 | d.169.1.8 | 1 | b.163.1.1 |

It is no surprise that most of these are from SCOP all-*β* class, as they are assemblies of *β*-strands, with class b.60.1.2, ‘Fatty acid binding protein-like’ as the exception. (However, because the *usage* option gives results for the entire `wwPDB`, the frequency may well reveal the experimental bias of solved structures within that family.)

Fig. SF2b shows the structural superposition of the regions from the linker region of the multimeric erythrocruorin from the earthworm, *Lumbricus terrestris,* (2GTL, chain o) and Human cellular retinol binding protein III (1GGL, chain a).

Are the erythrocruorin linker domains and the retinol-binding proteins homologues? Structural comparison shows that only the regions of these proteins share secondary structural elements and the geometry of their assembly (as shown in Fig. SF2b). The rest of the domains are quite different. The regions are not homologues. However, the dictionary has identified a substructure which they, and other domains, share. Like other supersecondary structures, their appearance in unrelated proteins show that they are shared pieces of protein folds but not signs of homology. Indeed, an appeal to SCOP shows that the erythrocruorin linker domains are in a superfamily of their own, sharing a folding topology with ‘Streptavidin-like’ domains.

**Fig. SF2.**
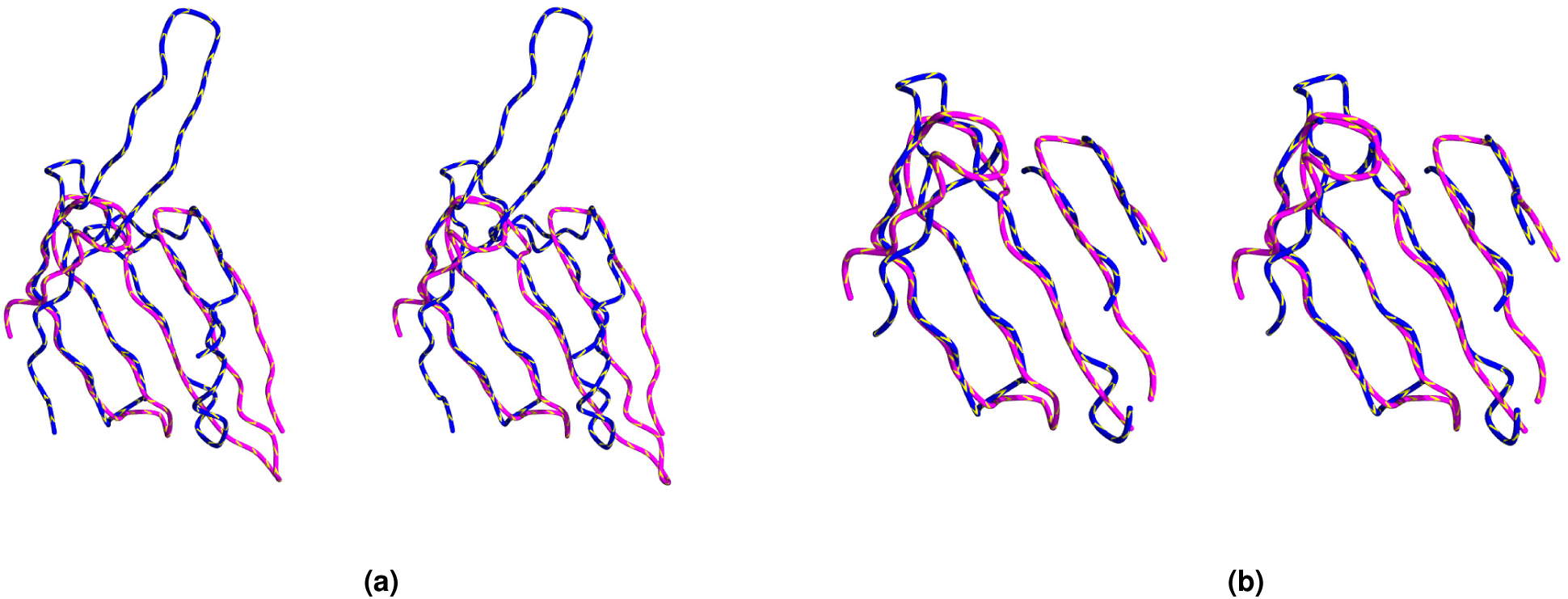
Superposition (in stereo) of region containing concept from (blue) linker region of earthworm multimeric erythrocruorin and (pink) Human cellular retinol binding protein III. (a) Entire region comprising concept `c_0559`. (b) Restriction to well-fitting residues from region comprising concept `c_0559`.

## Appendix

> The construction of Proçodic dictionary of concepts relies on the compression-based methodology we developed and presented at the 2017 IEEE Data Compression Conference (12). This methodology (with minor changes and additional details specific to this work) is reproduced below with permission for the convenience of the reader.

Our goal is to learn the static dictionary 𝒞 (i.e., hypothesis) that offers the best compression of the source collection 𝒯 (i.e, observed data). The general statistical framework used to achieve this relies on the criterion of minimum message length (MML) inference (17, 18). MML inference is best understood as a lossless two-part communication between an *imaginary transmitter-receiver pair*. In the first part the transmitter encodes and communicates the hypothesis to the receiver, while in the second part it communicates the observed data *given* the stated hypothesis. The best hypothesis in this framework is the one that yields the shortest two-part lossless message to communicate the observed data. Formally, for any static dictionary *𝒞* and source collection *𝒯,* the two-part message length (in bits) is denoted by the terms:

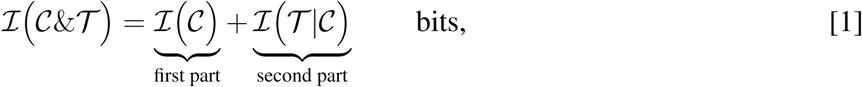

where ℐ(·) = −log_2_(Pr(·)) is the Shannon’s measure of information content (19).

The two-part message shown in Eqn. 1 is contrasted with the (single-part) *null model* message, that is, the encoding of the observed data *as is*, without the support of any hypothesis. The null model message length is denoted as ℐ _null_(*·*). Thus, the quality of an inferred dictionary *𝒞* is measured as the compression obtained by encoding the source collection *𝒯* using *𝒞*, i.e., as ℐ _null_ (*𝒯*) −ℐ(*𝒞*& *𝒯*) This yields an inference problem with the following objective: 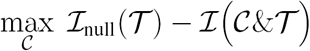

Addressing this inference problem requires the following: (1) A method to estimate the null model encoding length, ℐ _null_(*𝒯*), for any given collection *𝒯*; (2) A method to estimate the dictionary model encoding length, ℐ (𝒞 & *𝒯)*. for any given dictionary 𝒞 and collection *𝒯;* (3) A search method for an *optimal* dictionary (one that maximizes compression, as per the stated objective). These methods are presented below.

### Estimation of ℐ _null_(*𝒯*)

The null encoding of the source collection *𝒯* involves the encoding of the number of tableaux over an integer code, followed by the null encoding of each tableau *τ*_*𝓁*_ *∈𝒯* :

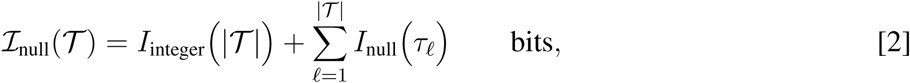

where *I*_integer_(.) is the message length of encoding any positive integer over a log*\** distribution (20). Further, the estimation of each *I*_null_ (*τ*_*𝓁*_) term is carried out by encoding the number of secondary structural elements using the same integer code, followed by encoding the three-tuples (**S**_*𝒯l*_ **Ω**_*τl*_, *Ξ*_*τl*_) using uniform probability distributions on their respective supports. This implies that each character in the S_*𝓁*_ string takes one bit to encode, each contact state in Ξ_*𝓁*_ also takes one bit, and each angle in Ω_*𝓁*_, specified to a precision of 0.1 ° in the range (−180 *°,* +180 *°*], takes log_2_(360*/*0.1) = log_2_ 3600 bits. Thus, the null message length for communicating each tableau *τ*_*𝓁*_ is given by:

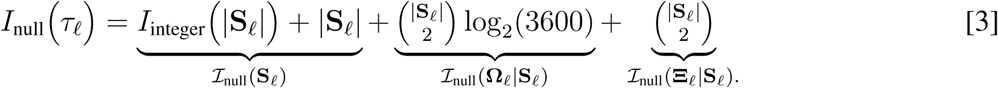

**Fig. SF3.**
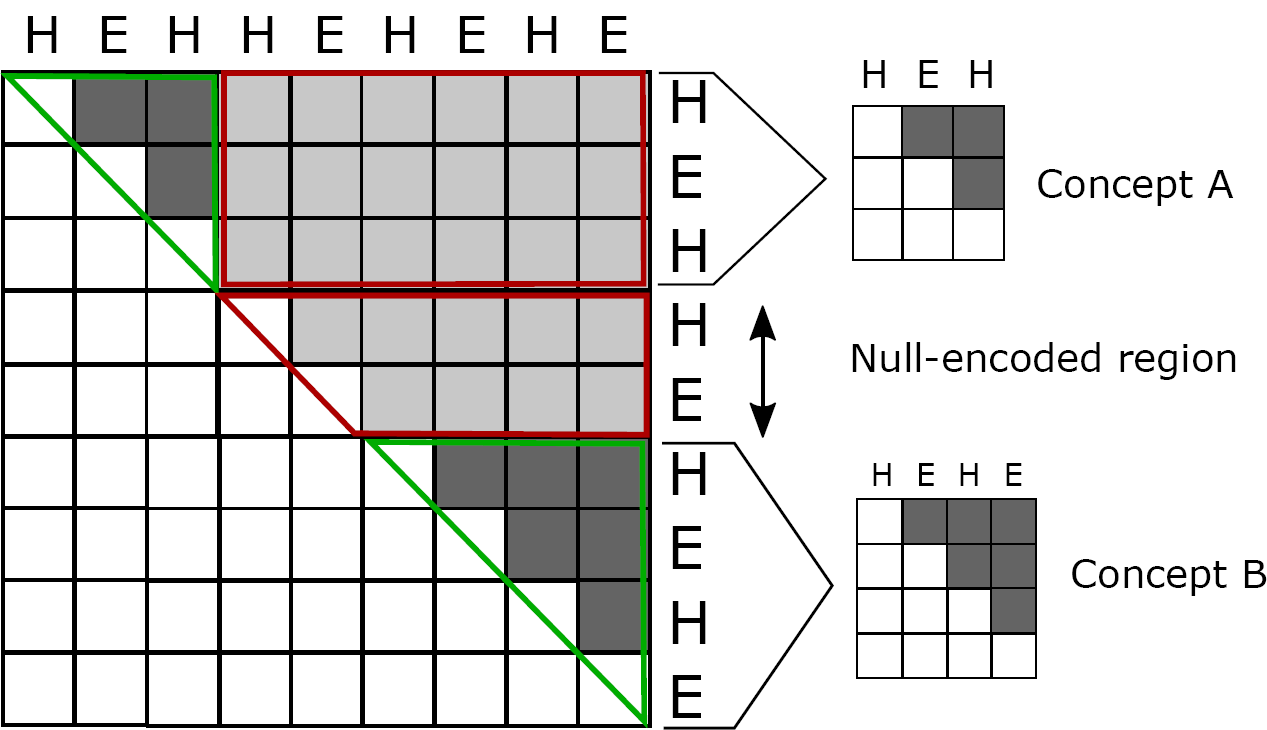
An illustration of a partition of a tableau.

### Estimation of the first part, ℐ(𝒞), term in Eqn. 1

Each concept *c*_*y*_ in any given dictionary is a (sub-)tableau. Therefore, *c*_*y*_ can be encoded using the null model as shown in Section S3, using **ℐ**_null_(*c*_*y*_) bits. In addition, its associated *κ*_*y*_ parameter also needs to be encoded. As seen in Section S1, each *κ*_*y*_ lies in the range [*κ*_min_, *κ*_max_] specified to a precision of *ϵ*_*κ*_, and can be encoded using a uniform probability distribution over this support. Using these component terms, the resulting encoding length for the full dictionary takes (in bits):

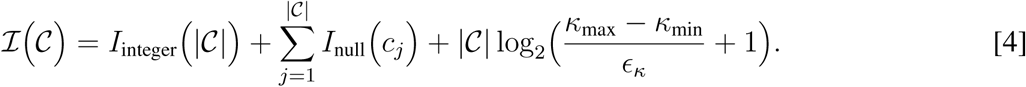

### Estimation of the second part, *ℐ* (*𝒯 | 𝒞*), term in Eqn. 1

The encoding length of the source collection *𝒯* given the dictionary *𝒞* is computed as the sum of the code lengths required to encode each tableau *τ*_*𝓁*_ *ϵ 𝒯* using 𝒞 To compute this code length,(*τ*_*𝓁*_ *| 𝒞*), each tableau *τ*_*𝓁*_ is *partitioned* into *non-overlapping* regions of variable sizes (see Fig. SF3). Any *partition* of *τ*_*𝓁*_, *p*(*τ*_*𝓁*_), is specified by an increasing sequence of integer indices 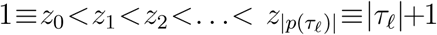. The set of possible 2-grams from this partition sequence, 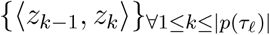, define |*p*(*τ*_*𝓁*_)| consecutive, non-overlapping regions of *τ*_*𝓁*_ of the form 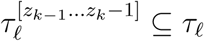. The total number of possible partitions for a given *τ 𝓁* is 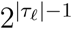

Assigned to each non-overlapping region 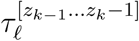 in a specified partition *p*(*τ* _*𝓁*_) is one of the concepts 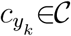 (provided 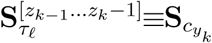), where 1≤*y*_*k*_ ≤|*𝒞*|, or a *null concept c*_0_. The key idea here is that the burden of explanation of the data in each non-overlapping region 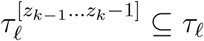 defined by *p*(*τ)* is borne by the corresponding data in its assigned concept. That is, the data in 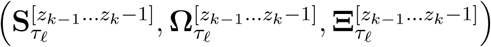 is communicated using the corresponding data 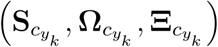 When the region is assigned to the null concept *c*_0_, the encoding of the data in that region is identical to null model encoding described in Section S3. In the remaining text, *p*(*τ*_*𝓁*_) specifies not only the non-overlapping regions, but also their corresponding concept assignments.

Thus, given *p*(*τ*_*𝓁*_), the data within each *τ*_*𝓁*_ ∈ *𝒯* can be communicated as follows. First, the tableau size is communicated with an integer code using *I*_integer_ (|*τ*_*𝓁*_ |) bits. Second, the details specified by *p* (*τ*_*𝓁*_) are communicated using *ℐ* (*p*(*τ*_*𝓁*_))bits. Third, each non-overlapping region 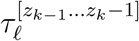 is explained by its assigned concept 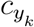 (shown as the grey coloured regions in Fig. SF3), using 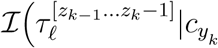 bits. Finally, the remaining data of *τ*_*𝓁*_ (the light-grey cells in Fig. SF3) are communicated using the null model (Section S3). We denote this data (using set notations) as 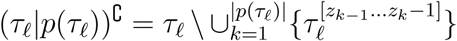, with code length denoted by *ℐ*_null_((*τ*_*𝓁*_ |*p*(*τ*_*𝓁*_))`C`).

Based on the above details, the *shortest* lossless encoding length of any tableau *τ*_*𝓁*_ *∈ 𝒯* given a static dictionary *𝒞*, is one that that minimizes the following objective:

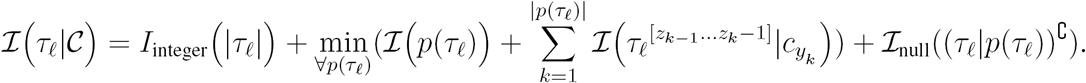

Combining the above term with Eqn. 4, gives us *ℐ* (*𝒯* | *𝒞*)as:

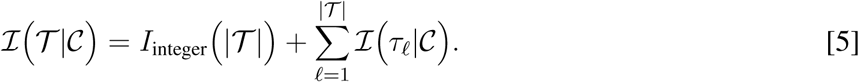

Below we describe the details of computing the code length terms involved in Eqn. 5.

### Computation of *ℐ* (*p*(*τ*_*𝓁*_))

A partition *p*(*τ*_*𝓁*_) is encoded as follows. Since the tableau size |*τ*_*𝓁*_ | has already been communicated, the size of the partition |*p*(*τ*_*𝓁*_)| is encoded in log_2_ |*τ*_*𝓁*_ | bits. Each index in the corresponding set of concept assignments 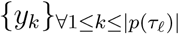 can take the values 0 ≤ *y*_*k*_ ≤ | *𝒞* |, and, thus, can be encoded in log_2_(|*C*| +1) bits. Given this information, the values of 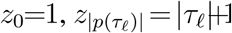, and the subset of {*z*_*k*_}’s (2 *≤ k <* |*p*(*τ*_*𝓁*_)|) associated with regions *not* assigned to the null concept *c*_0_ are already decipherable based on the assigned concept sizes. The remaining ones, associated with *c*_0_, each take log_2_(|*τ*_*𝓁*_ |*-z*_*k-*1_+1) bits to state.

### Computation of 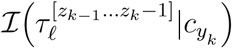

A region 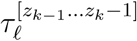 can be encoded using a null concept *c*_0_, or using any concept 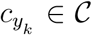. The computation of the null concept encoding of a region follows the same scheme as the null-model encoding of a tableau described in Section S3. On the other hand, the encoding of 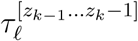 using a concept 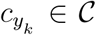 is permitted only when the corresponding secondary structural strings are identical, and when the corresponding contact information between pairs of secondary structural elements differ in no more than 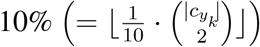 places.

The details of encoding 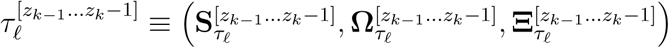 with 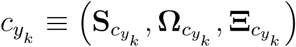 are now considered. Since 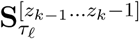 is identical to 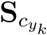, it is only necessary to encode 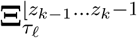 and 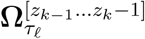 using the corresponding concept’s matrices 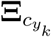 and 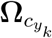, respectively. Thus, the encoding *N*_*m*_ requires 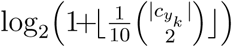 bits, where 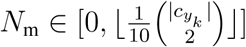 is the total number of mismatches in this assigned region where 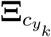 and 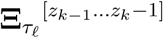 differ. The locations of the mismatches are then encoded using a uniform distribution over the number of ways of identifying *N*_*m*_ locations out of 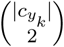 cells. The resulting code length to identify each mismatched entry takes the logarithm of the corresponding binomial coefficient. Once this information is communicated, the corresponding entries of 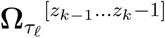 are encoded using the null model, each using log_2_ 3600 bits.

The matched angles are now transmitted. Each signed angle 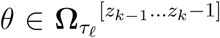 is encoded using the corresponding angle 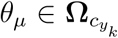 and a (90%, 10%) mixture model of von-Mises circular distribution (18) (parameterized on *θ*_*µ*_ and concept’s *κ*) and a uniform distribution over the support (*-*180°, +180°].

### Computation ℐ_null_((*𝒯*_*l*_*|p*(*𝒯* _*l*_))^C^)

This refers to the encoding of the off-diagonal 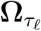 and 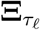 entries, shown as the light-grey coloured cells in Fig. SF3, and denoted here as 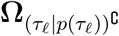 and 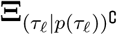. The total number of entries in this off-diagonal area in each of these matrices is 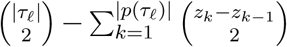. Each angle in 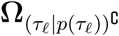 is encoded using the null model in log_2_(3600) bits. Each contact in 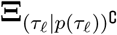 requires 1 bit.

### Optimal partition of *τ*_*𝓁*_ given *𝒞* via dynamic programming

The computation of *ℐ(τ*_*𝓁*_ *|𝒞)* is carried out on the optimal partition of *τ*_*𝓁*_ given the concepts in *𝒞*. The identification of the optimal *p*(*τ*_*𝓁*_), the one that minimizes *τ*_*𝓁*_, is achieved using a one-dimensional dynamic programming algorithm, similar to the one devised in our earlier work for a *completely different* problem from the same domain (11).

To compute the optimal partition *p*(*τ*_*𝓁*_) of a tableau *τ*_*𝓁*_ under the dictionary *𝒞*, we first construct a set {*H*^0^,*H*^1^,…,*H*^|*𝒞|*^} of *|𝒞|* + 1 matrices called the “code length matrices”. The (*i, j*)-th entry of code-length matrix *H*^*t*^ stores the message length for stating the subtableau region 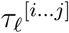 using the concept *c*_*t*_*∈ 𝒞* plus the corresponding off-diagonal entries in that region 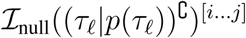. (If the region is stated using the null concept, this entry also adds the cost of stating the size of the null region.)

The manner in which partitioning has been formulated allows us to investigate the problem of determining the optimal partition by solving smaller independent sub-problems, thereby allowing us to propose a dynamic-programming (DP) approach. We first consider a cost function *M* (*j*) defined using the following recurrence relationship, for every integer *j* such that 0 *≤ j ≤ |τ*_*𝓁*_*|* :

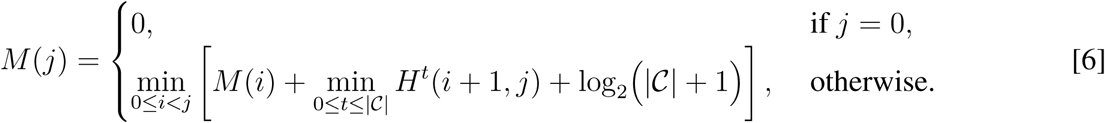

The recurrence relation in Eqn 6 implies that *M* (*j*) is the optimal cost of stating the *τ*_*𝓁*_ up to the SSE index *j*. This can be verified as follows. For *j* = 0, *M* (*j*) is interpreted as the cost of stating nothing, which is zero. For any other value *j,* we find the optimal dissection point *i < j* such that the tableau up to SSE *j* can be stated incrementally as the tableau up to SSE *i* using some dissection, plus the rest of the tableau up to SSE *j* using a concept that best explains the region [i … j]. We can hence compute the optimal statement cost of the dissection of the entire tableau, *M* (*j* = *|τ*_*𝓁*_*|*). Adding to it the cost of stating the number of SSEs in the tableau and the cost of stating the number of segments in the optimal segmentation, we will have the expansion for *ℐ(τ*_*𝓁*_*| 𝒞*) term in Eqn. 5 as:

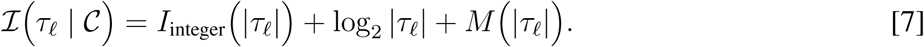

The optimal segmentation can then be determined by tracing back on the cached optimal derivations of *M,* starting from *M* (*j* = *|τ*_*𝓁*_*|*) back to *M* (*j* = 0).

### Searching for an optimal dictionary

Our goal is to address the problem of inferring an optimal static dictionary, i.e., one that minimizes the two-part message length given by Eqn. 1. Finding a provably optimal dictionary is computationally intractable due to the enormous search space. Hence, a simulated annealing (SA) heuristic is devised to address this problem. Algorithms based on SA require an aperiodic irreducible Markov chain defined on a certain state space, and a *cooling schedule* to push iteratively the solution towards the optimum. In our case, the state space is the set of all possible dictionaries. The desired Markov chain is generated by defining a neighbourhood and the corresponding transition probabilities for every state 𝒞. A local neighbourhood for every state is explored through the following perturbation primitives: (1) `Add concept:` Creates a concept randomly from the source collection and adds it to 𝒞. (2) `Remove element:` Chooses a concept randomly from the dictionary and deletes it. (3) `Perturb concept length:` Chooses a concept randomly from the dictionary, and extends/shortens it, in reference to its original source. (4) `Perturb concept kappa:` Increments/decrements the current value of *κ* associated with a randomly chosen concept. (5) `Swap concept with usage:` Chooses a concept randomly from the dictionary, and swaps it with a region in the collection that is currently encoded by it. (Note, the usage swapped-in as the perturbed concept could weakly violate the connectivity constraint in terms of its contact map, unless strict connectivity is also imposed on that chosen usage.)

At each iteration, one of the above five perturbations is chosen uniformly at random. The transition probability to the neighbour is computed as follows. If the two-part message length given by Eqn. 1 decreases, the transition probability to the perturbed state is 1. If the two-part message length increases by Δ*I* bits, the probability is 2^*-*Δ*I/T*^, where *T* is the *temperature* parameter of the system controlled by the following cooling schedule: Start with a temperature of 5,000 and decrease it by a factor of 0.88. At each temperature step, perform 50,000 random perturbations unless the temperature is below 10, where the number of perturbations is increased to 500,000 per temperature step. When the temperature reaches below 0.1, the search stops and the current state of the dictionary is reported. See `pseudocode.pdf` (click).

### Parallel Implementation

The methodology of inference described above was implemented in the `C++` programming language. To tackle the enormous amount of computational effort that was required to identify the Proçodic dictionary reported here, this program was parallelised and deployed on a large computing cluster. This was achieved using the `OpenMPI C++` library, under a Message-Passing Interface framework.

Specifically, this implementation was parallelized and was executed over 240 computing cores (requiring about 2 Giga Bytes of main memory per core) on the high performance computing cluster, `Tinaroo`, at the University of Queensland’s Research Computer Center. It took roughly 11 days for our simulated annealing approach to converge on the identified Proçodic dictionary of 1493 concepts – this runtime is tantamount to about 7 years of computing over a sequential run.

The source collection gets sub-divided into roughly equal subsets of tableaux, and allocated to individual computing nodes. This data-parallel implementation exploits the ability for each computing node to generate *independently* the same sequence of uniform (pseudo-)random numbers. At each node, and operating on the allocated subset of tableaux from the collection that is being compressed, all random numbers required for performing the perturbations involved in the simulated annealing approach are ensured to work in tandem across the cores. This results in every node evolving the same dictionary independently, although working on different partitions of the source collection. This minimises the overhead required for communication between the compute nodes, by avoiding the overhead involved in broadcasting the current state of the dictionary and the details of the perturbations being performed at each perturbation step of the simulated annealing approach. The only communication required between nodes is the ‘*Allreduced*’ total two-part message length, *summed* over all data partitions, before perturbations are synchronously accepted/rejected (as per the Metropolis criterion). The total speedup as a result of this parallelisation was measured to about 200 times over 240 nodes.

Conceptualization: ASK; Methodology: AML, LA, DA, PJS, MG, and ASK; Software: RS and ASK; Validation: AML, LA, and ASK; Analysis: AML, LA and ASK; Investigation: AML, RS, PJS, MG and ASK; Resources: AML and DA; Data Curation: ASK; Writing - Orginal Draft: AML and ASK; Writing - Review & Editing: RS, LA, DA, PJS and MG; Visualization: AML and ASK; Supervision: ASK; Project Administration: ASK; Funding Acquisition: AML, PJS, MG and ASK.

